# FrlP, an ABC type I importer component of *Bacillus subtilis*: regulation and impact in bacterial fitness

**DOI:** 10.1101/2025.07.11.664333

**Authors:** Inês C. Gonçalves, Ana Pontes, Carla Gonçalves, Isabel de Sá Nogueira

**Affiliations:** Microbial Genetics Laboratory, UCIBIO, Department of Life Sciences, NOVA School of Science and Technology, Universidade NOVA de Lisboa, Caparica, Portugal; Yeast Genomics Laboratory, UCIBIO, Department of Life Sciences, NOVA School of Science and Technology, Universidade NOVA de Lisboa, Caparica, Portugal; Associate Laboratory i4HB, NOVA School of Science and Technology, Universidade NOVA de Lisboa, Caparica, Portugal

**Author notes:** Address correspondence to Isabel de Sá-Nogueira.

## Abstract

*Bacillus subtilis* is able to catabolize fructosamines, also known as Amadori rearrangement products. The *frlBONMD-frlP* operon mediates this process and is subjected to specific and global regulation. Although the degradation pathway favoring α-glycated amino acids is known, the mechanisms of substrate uptake have remained unclear. In this study, mutagenic and functional analyses revealed that FrlONM, a type I ABC importer, along with the nucleotide-binding domain (NBD) FrlP, is required for the uptake of fructosevaline. Transcriptional and translation *frlP-lacZ* fusions indicated that *frlP* is induced by fructosevaline and negatively regulated by the FrlR repressor. Furthermore, by targeting an RNA molecule (S1254) encoded upstream from *frlP* and overlapping the repressor *flrR* gene transcribed in the opposite direction, we provide evidence for the dual regulation of *frlP* by this anti-sense RNA and FrlR. This result adds a posttranscriptional layer to the intricate regulation of the fructosamines operon. In addition, we show that MsmX, a multitask NBD of *B. subtilis*, is also able to serve as energy motor of this type I ABC importer, and that its presence alongside FrlP is vital for optimal growth on fructosevaline. To address the physiological significance of this functional redundancy we assessed the distribution of ABC type I NBDs FrlP and MsmX across the *Bacillaceae* family. MsmX is homogeneously distributed in *Bacillaceae* family tree, while FrlP is restricted to the *Bacillus subtilis* group, suggesting that the presence of FrlP together with other components of the fructosamines operon is important for bacteria fitness in plant-associated ecological niches.

**Importance:** *Bacillus subtilis* is widely applied in the industry as a microbial cell factory, as a biofertilizer for sustainable agriculture, in the animal feed industry and as human probiotic. In its natural environment *B. subtilis* helps to shape the gut microbiome and the phytomicrobiome. Fructosamines, or Amadori rearrangement products, are ubiquitously found in nature and serve as precursors of toxic cell end-products implicated in the pathology of human diseases. This study provides a solid contribution to a deep knowledge of transport mechanisms, genetic regulation and physiological relevance of fructosamines utilization in *B. subtilis*. Moreover, it highlights an unusual strategy to adapt to alterations in nutrient availability by swapping the energy providing domain of ABC transporters.

## Introduction

Ubiquitous across all kingdoms of life, ATP-binding cassette (ABC) transporters share a similar structural architecture which consists of two transmembrane domains (TMDs), that assemble the translocation pore enclosed in the lipid bilayer, and two nucleotide binding domains (NBDs or ATPases) that energize the transport system via ATP hydrolysis (1–4). Contrarily to TMDs, NBDs share sequence and folding conservation, especially motifs responsible for accommodation and coupling of the ATPase activity (4). ABC exporters are transversal to all domains of life, whilst importers are exclusive to prokaryotes and plants (5). ABC Type-I importers are accountable for the transport of primary metabolites needed in high quantities, such as sugars and amino acids, and they require a third component, a solute-binding protein (SBP), which in Gram-negative bacteria is located in the periplasmic space while in Gram-positive organisms is anchored to the plasma membrane through a N-terminal extension (4, 6). In bacterial genomes, all elements of ABC transporters are generally encoded in the same DNA region or clustered together in an operon (7, 8). That is the case of the prototype of bacterial ABC Type-I importers the MalEFGK_2_ from *Escherichia coli* responsible for maltose uptake (8, 9).

The chromosome of the well-studied Gram-positive bacterium *Bacillus subtilis* revealed exceptions displaying several independent but incomplete ABC carbohydrate transport systems lacking the NBD and two orphan NBDs, MsmX and FrlP (formerly YurJ) (7). Studies have shown that the orphan ATPase MsmX is able to energize six different carbohydrate uptake systems, namely MdxEFG (10), AraNPQ (11), GanSPQ (12), RhiLFG and YtcPQ-YteP (12, 13), and MelECD (14), involved in the uptake of maltodextrins, arabino-oligosaccharides, galacto-oligosaccharides, galacturonic acid oligomers and/or rhamnose-galacturonic acid disaccharides and melibiose, respectively. Due to the extended capacity of interaction and energization of a large spectra of transporters, MsmX was denominated as multitask or multipurpose (11). The less studied FrlP ATPase is encoded in the downstream region of an operon responsible for fructosamine utilization (Fig. 1) (15, 16). This operon comprises *frlB*, encoding a deglycating protein, *frlD*, encoding a kinase, putative transporter genes (*frlONM*) and by a convergently encoded repressor gene, *frlR*, co-localized with a non-coding RNA, S1254 (15, 16).

**Fig. 1.**
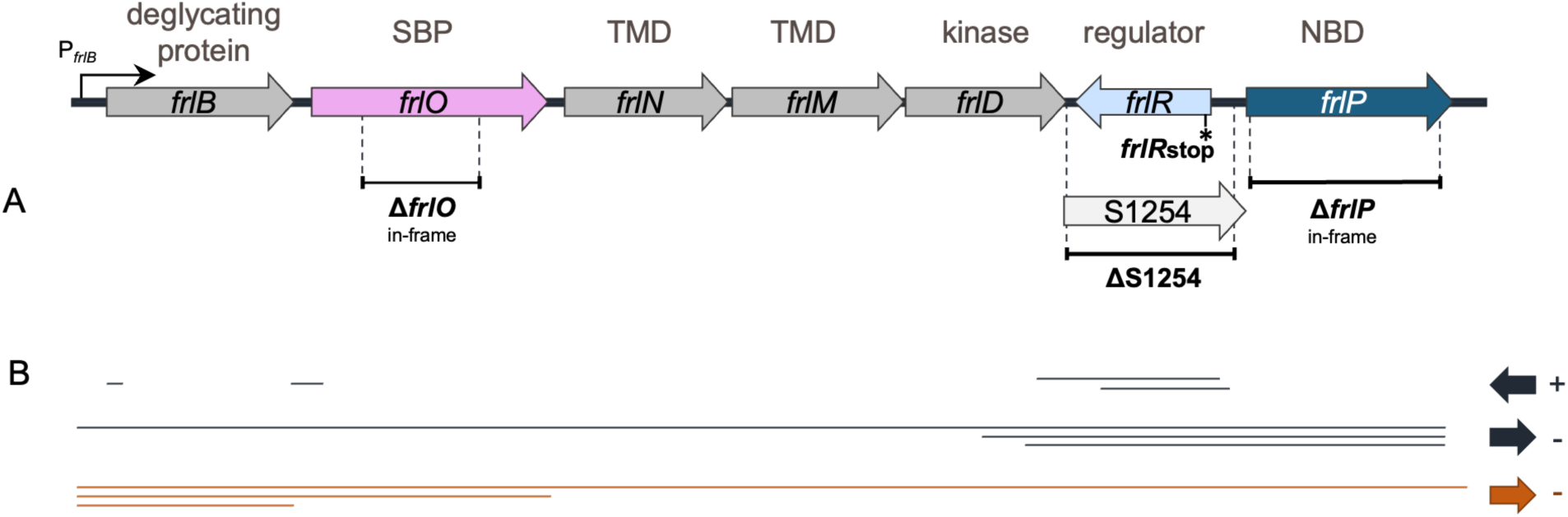
Representation of the *frlBONMD* operon genomic region enclosing genes coding for the glycosidase FrlB, the substrate binding protein (SBP) FrlO, permeases (TMDs) FrlN and FrlM, the kinase FrlD, the repressor of the operon, FrlR, and the nucleotide binding domain (NBD) FrlP. The putative antisense RNA S1254 is located between *frlD* and *frlP* sequences. The promoter of the operon P*_frlB_ is* indicated by an arrow. **(A)** Schematic representation of *frl* operon mutations constructed in this work: in-frame deletion Δ*frlO*; nonsense point mutation *frlRstop*; deletion ΔS1254; and *in-frame* deletion Δ*frlP*. **(B)** Transcription units identified by ChIP-chip experiments (17), retrieved from SubtiWiki (http://www.subtiwiki.uni-goettingen.de), are represented below as solid dark lines and arrows point to the direction of transcription. Solid orange lines indicate mRNA molecules detected by northern blot analysis using a *frlB* probe after cell growth in minimal medium with glucose (30).

Fructosamines, or Amadori products, constitute the first stable intermediates of the Maillard reaction, being spontaneously produced when the carbonyl group of reducing sugars non-enzymatically reacts to the free amine group of amino acids or proteins (18). Amadori products are ubiquitously found in nature, such as humic substrates, fruits and vegetables, and in several stored or processed foods (19, 20). In certain ecological niches, for example the human and animal gut or the rhizosphere, Amadori products shape the microbiome (19, 21, 22). *E. coli*, yeasts and mammals are subjected to intracellular glycation, so it is plausible that the same could happen in *B. subtilis*, driven by the presence of glucose 6-phosphate(P), resulting from the metabolism of other carbon sources (15). Microorganisms can metabolize fructosamines through deglycation by two types of enzymes: amadoriases (oxidases) or fructosamine kinases, the latter being the mechanism observed in both *E. coli* and *B. subtilis* (15, 16, 23). Fructosamine kinases phosphorylate the amadori product at C-6 and the deglycase enzyme catalyzes the beakdown of the fructosamine 6-P into *e.g.* glucose 6-P and a free amine (15, 23, 24). Furthermore, it has been shown that both organisms are able to sustain growth on Amadori products (16, 23). However, substrate specificity differs between them; unlike *E. coli*, *B. subtilis* kinase FrlD and deglycase FrlB have substrate specificity towards α-glycated amino acids instead of ε-glycated lysine (24).

Although some efforts have been made to understand the physiological significance of the *frlBONMD* gene products in *B. subtilis* (16, 24), targeted gene studies are still lacking, especially regarding *frlP* and its involvement in this fructosamine utilization pathway. Our previous studies have shown that FrlP is able to energize AraNPQ and GanSPQ in the import of arabinan and galactan derived products, respectively, upon ectopic expression controlled by a synthetic promoter (12, 25). Here we report by genetic and functional studies the involvement of FrlONM-FrlP in the uptake of fructosevaline. Moreover, by genetic analysis it is revealed the possible involvement of MsmX in this sugar amines utilization pathway. Transcriptional and translational studies of *frlP* expression in the presence of different substrates disclosed a dual mechanism of regulation and shed light on the putative exchangeability between FrlP and MsmX and its role in bacterium fitness. The importance of FrlP presence in the cell was further investigated by exploring the phylogenetic distribution of both FrlP and MsmX amongst the *Bacillaceae* family.

## Results

### FrlONM are involved in the uptake of fructose valine

The *frlBONMD* operon (Fig. 1) has previously been reported as necessary for the uptake and utilization of fructosamines based on studies using a *frlBONMD* knock-out mutant (16). FrlP has been proposed as the ATPase that couples the transport of fructosamines (26–28), even though its functional role has not been experimentally validated. Similarly, FrlO has been annotated as the SBP responsible for Amadori delivery to TMDs FrlN and FrlM (27, 29). However, previous studies used a raw mixture of the Amadori product fructosyl-arginine that retained residual glucose and arginine (16) or small amounts of product obtained by an undisclosed protocol (28). Since *B. subtilis* kinase FrlD and deglycase FrlB have substrate specificity towards α-glycated amino acids (24) in this study we used fructosevaline (mixture of diastereomers; CymitQuimica). To ascertain the role of FrlONM in the import of fructosamines, a markerless *in-frame frlO* deletion mutant was constructed to prevent polar effects on the transcription of the operon downstream genes (Fig. 1A). The ability of the mutant strain to sustain growth with fructosevaline as sole carbon and nitrogen source was determined and compared to the WT strain (Fig. 2). The results showed that Δ*frlO* mutant failed to grow on fructosevaline, confirming that FrlO is essential for the delivery of this Amadori product to the TMD domains of the FrlONM transporter complex.

**Fig. 2.**
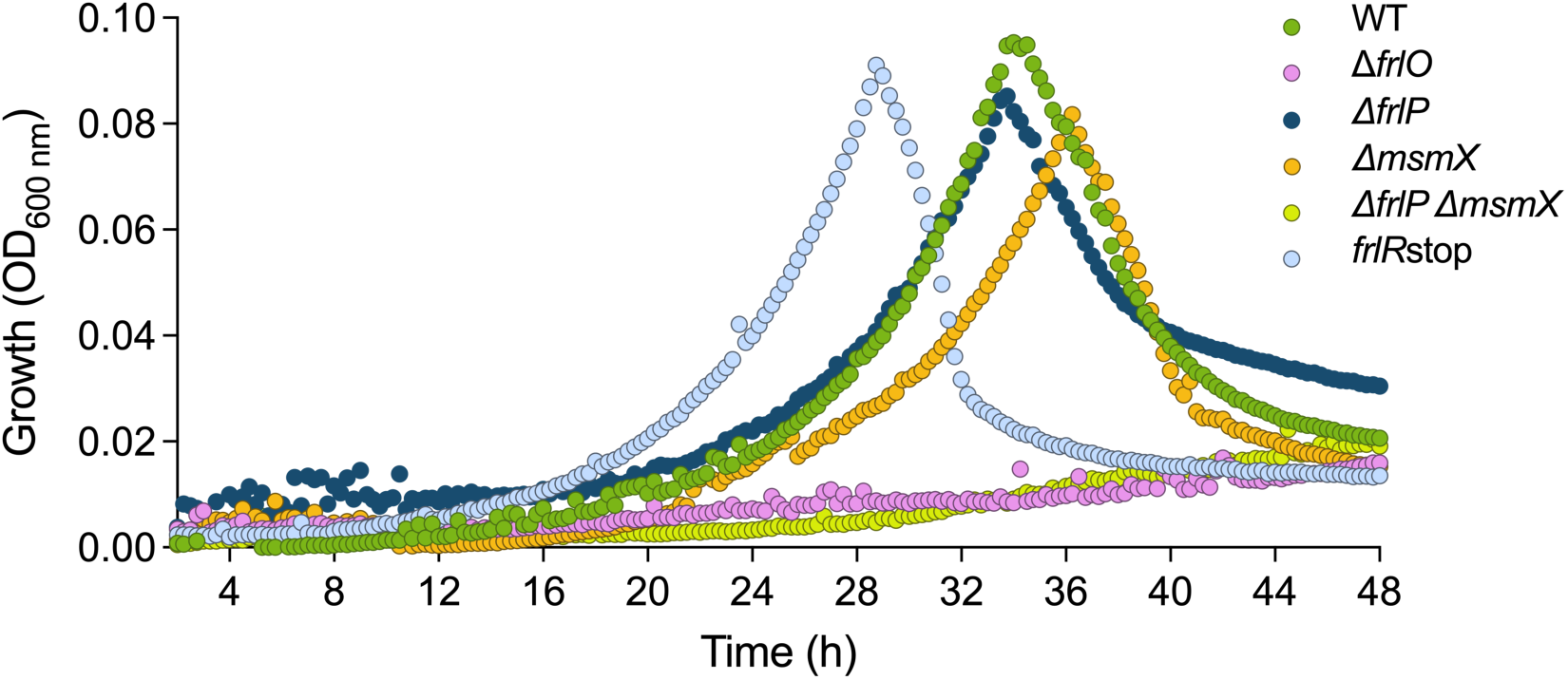
Growth curves of *B. subtilis* WT (dark green), *frlRstop* (light blue), Δ*frlO* (pink), Δ*msmX* (orange), Δ*frlP* (dark blue) and Δ*msmX*Δ*frlP* (light green) strains in M9 minimal medium supplemented with 22 µg mL^-1^ CAF and 2 mM fructosevaline. Shown is one representative growth curve of each strain. Two independent growth experiments were performed using technical triplicates, except for the Δ*frlP*, for which the four independent experiments were conducted. All growth curves are shown in Supplementary Fig. 1C.

### FrlP and MsmX are two NBDs that energize the FrlONM importer

The *frlP* gene encoding a putative NBD domain of the FrlONM was previously shown to be part of the *frlBONMD* operon by tiling arrays and northern blot (17, 30; Fig. 1B). To determine if FrlP is responsible for energizing the FrlONM transport system, a deletion mutation Δ*frlP* was constructed and the resulting strain was assessed for its ability to grow on fructosevaline as the sole carbon and energy source (Fig. 2). The deletion did not impair the ability of the mutant Δ*frlP* cells to grow on fructosevaline, suggesting that other NBD(s) might energize the FrlONM transport system. Previously, our laboratory showed that ectopic expression of *frlP*, under the control of a synthetic promoter P_spank(hy)_, complements the absence of the multitask NBD MsmX in the oligosaccharides transport systems AraNPQ and GanSPQ (25). Thus, we tested the ability of an *msmX* null mutant (Δ*msmX*) to grow on fructosevaline as the sole carbon and energy source. Remarkably, the impact of the *msmX* deletion on bacterial growth was very similar to that displayed by the *frlP* deletion (Fig. 2). We therefore generated a *msmX* and *frlP* double mutant which showed to be unable to grow on fructosevaline (Fig. 2). These results strongly suggest that neither FrlP nor MsmX function as the exclusive energy generator components of the fructosamines transporter complex. Under the conditions tested, both NBDs (FrlP and MsmX) are functionally interchangeable and capable of energizing the FrlONM transporter.

To confirm this line of evidence, protein-protein interactions of MsmX and FrlP with TMDs FrlN and FrlM were probed by a bacterial two-hybrid system (B2H) in *Escherichia coli* (31). For validation, the interaction between MsmX and AraQ, a TMD from the AraNPQ importer, was also tested. The results revealed that both FrlP and MsmX can establish *in vivo* contacts with TMDs FrlN and FrlM, corroborating their functional interchangeability as NBD domain of the importer FrlONM (Fig. 3).

**Fig. 3.**
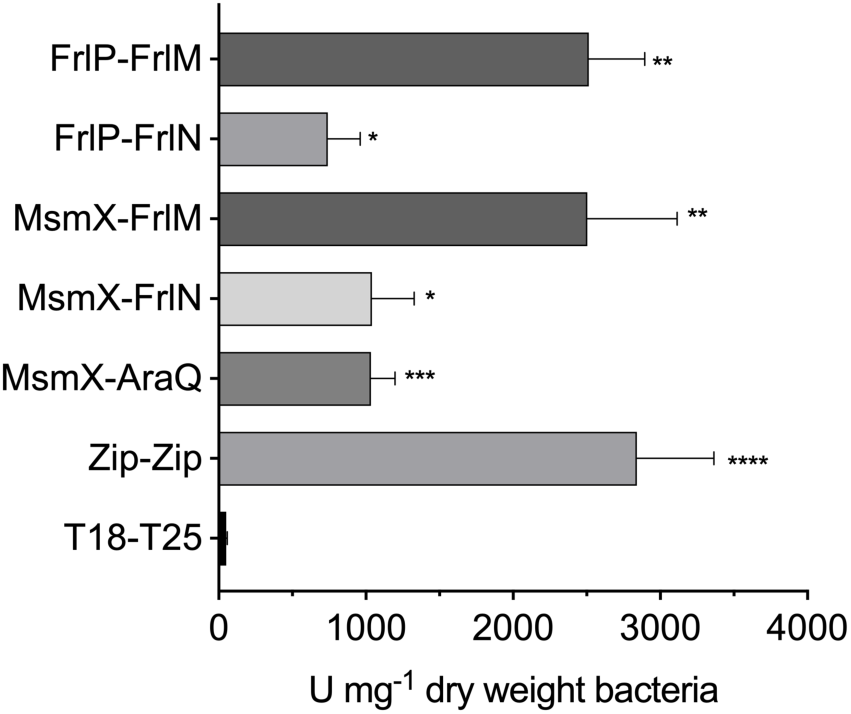
Quantitative analysis of protein-protein interaction by B2H assays. Interaction between T18 and T25 fragments expressed from empty pUT18 and pKT25 vectors was used as negative control. Zip-Zip and MsmX-AraQ interactions were used as positive controls. At least three independent experiments are represented for each interaction; the error bars represent the standard deviation of the mean. *β*-galactosidase activity was calculated as described in Material and Methods and it is represented in U mg^-1^ dry weight bacteria. Statistical significance between *β*-galactosidase activity of the different interactions and the negative control is indicated (*p<0.05; **p<0.01; ***p=0.001; ****p<0.0001).

### Expression of *frlP* is induced by fructosevaline and negatively regulated by FrlR

Repressors CodY and FrlR were previously shown to bind directly to the *frlB* promoter region, controlling the expression of the fructosamine operon (16, 32, 33). More recent studies contradict these observations proposing that FrlR functions as an activator (28). To clarify the controversy and analyze the regulation of *frlP* we constructed a nonsense mutation at the 5’-end of the gene (*frlRstop*; Fig. 1A). When growing this strain in the presence of fructosevaline the mutant reached its peak of growth earlier than the WT strain (Fig. 2 and Fig. S1C) suggesting that FrlR acts as a repressor. This behavior is identical to that observed with the repressor of the ε-fructoselysine operon of *E. coli* (34).

Expression of *frlP* is driven by the promoter P*_frlB_* (17, 30; Fig. 1A), however by bioinformatic analyses a putative promoter was found upstream from the *frlP* coding region (17). In addition, although FrlP can functionally substitute MsmX in the import of arabinotriose when ectopically expressed, the expression in its own *locus* does not lead to protein accumulation in the presence of glucose, arabinose or arabinotriose as sole carbon and energy source (12). To answer these questions and understand how *frlP* is expressed in its own locus, transcriptional and translational fusions of the *frlP* 5’-end to the *lacZ* gene of *E. coli* were constructed (Fig. 4A). The fusions were analyzed in both WT and *frlRstop* nonsense mutant backgrounds grown in the presence of different carbon sources: fructosevaline, glucose, fructose and casaminoacids (Fig. 4B). *β*-Galactosidase activity assays showed very low level of *frlP* expression in the WT background when grown in glucose, fructose or casamino acids, both at transcriptional (T*_C_*) and translational (T*_L_*) levels, whereas a high increase of about 150-fold (T*_C_*) and 225-fold (T*_L_*) in the expression was observed in the presence of fructosevaline when compared to fructose. This result suggests that fructosevaline, or one of its metabolic products, is a strong inducer of *frlP* expression. In the depleted FrlR mutant (*frlRstop*), *frlP* expression increased about 18-fold (T*_C_* and T*_L_*) in the presence of glucose and 40 (T*_C_*) to 70-fold (T*_L_*) in the presence of fructose, further indicating that FrlR acts as a repressor, impairing transcription and translation in the presence of these sugars. In the presence of casamino acids repression is only partially relieved, which is consistent with the role of CodY, as a negative regulator of the fructosamines operon (16, 32, 33). In sum, our data correlate with previous studies conducted with the promoter P*_frlB_* (16), indicating that *frlP* expression is induced by fructosevaline and negatively regulated by FrlR and CodY.

**Fig. 4.**
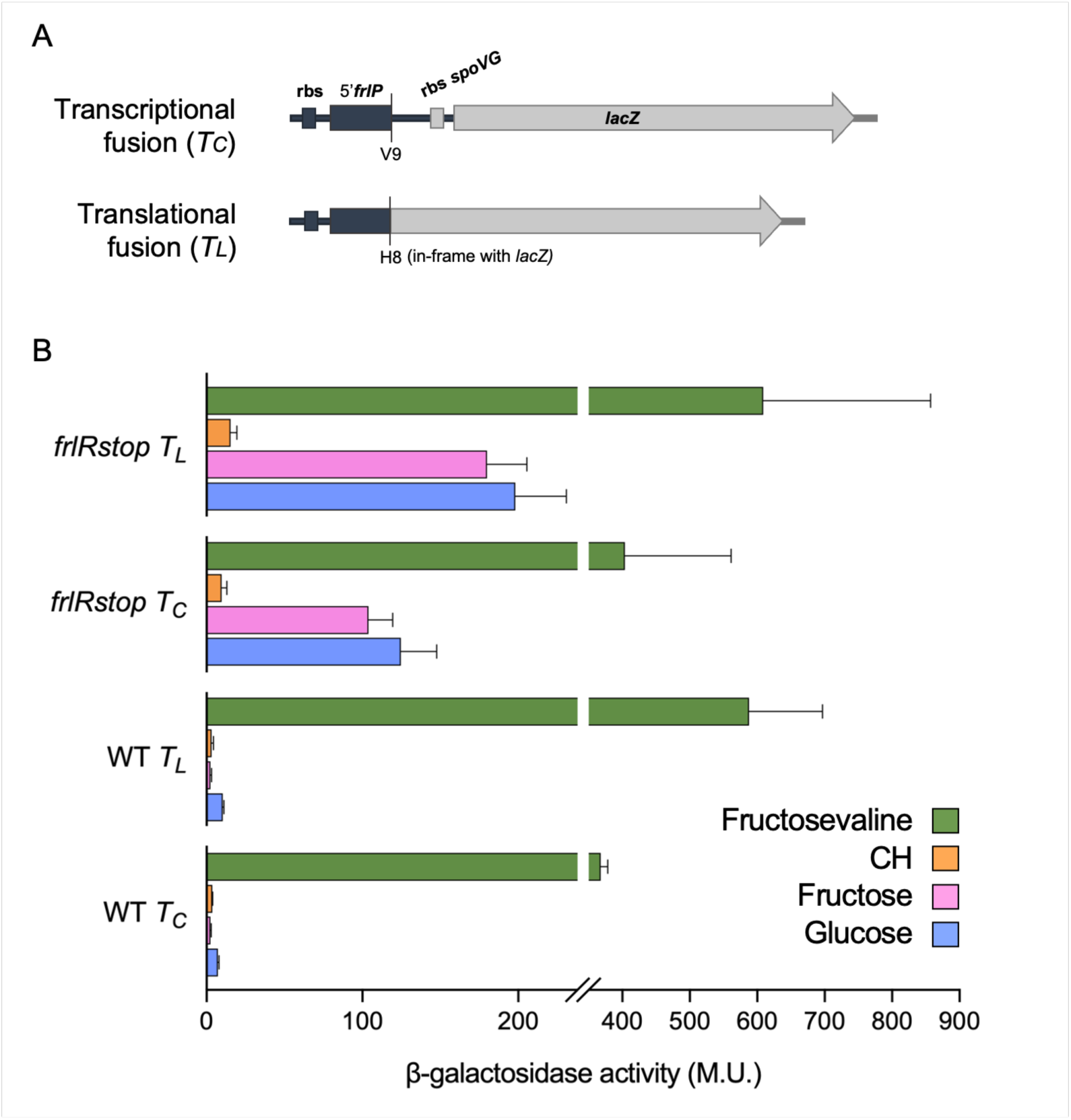
Transcriptional (T*_C_*) and translational (T*_L_*) gene fusions constructed for measurements of *frlP* expression. **(A)** Genomic representation of transcriptional and translational fusions of the 5’-end region of *frlP* gene to the *lacZ* gene from *E. coli*. **(B)** Quantification of *frlP* expression in the WT and *frlRstop* genetic backgrounds. *β*-galactosidase activity of 5’*frlP*-*lacZ* strains grown in M9 minimal medium supplemented with 22 µg mL^-1^ CAF and 1 mM glucose, 1mM fructose, 1 % (wt/vol) casamino acids (CH) or 2 mM fructosevaline, was measured in cells recovered at approximate growth peaks (Fig. S2). M. U. – Miller Units. At least two biological replicates are represented. In glucose, *β*-galactosidase activity of T*_L_* fusion in the WT background (strain ISN129) was measured in two independent experiments, while expression data from the remaining strains include at least three independent experiments. Assays in fructose and CH were measured three and two times, respectively, for all strains. *β*-galactosidase activity in fructosevaline was repeated two times, except for the WT strain, for which three independent experiments are included.

### Accumulation of FrlP is under the dual control of FrlR and S1254 RNA

FrlP can functionally substitute MsmX at energizing the AraNPQ transporter when expressed ectopically (12). However, previous studies failed to detect accumulation of FrlP when expressed at its own *locus*, although mRNA levels of *frlP* are high, pointing to the existence of a genomic context-dependent post-transcriptional mechanism involved in the regulation of *frlP* expression (12). The gene *frlP* is the last gene of the transcriptional unit *frlBONMD*-*frlP*, and it is co-transcribed with a putative antisense RNA S1254 overlapping the entire regulator gene *flrR*, which is transcribed in the opposite direction (Fig. 1)(16, 17). Thus, we postulated that either S1254 and/or the repressor of the operon, FrlR, could be involved in the regulation of *frlP*. To test this hypothesis, we constructed a markerless deletion of S1254 in a *msmX*-null background. Since this deletion also encompasses the overlapping regulator gene *frlR*, an additional strain was constructed carrying a *frlR* nonsense mutation (*frlRstop*) in a *msmX*-null (Δ*msmX*) background. In this mutant background, the entire operon transcription unit is maintained intact, but the point mutation impairs the production of the repressor FrlR (Fig.1). These strains were functionally assessed by their ability to substitute MsmX as the energizer of the AraNPQ import system. Growth kinetics parameters were measured in the presence of the substrate of the importer, arabinotriose, as sole carbon and energy source and the results were compared to the WT, single Δ*msmX* mutant, and the strain ectopically expressing *frlP* under the control of a synthetic promoter P_spank-hy_. The findings are summarized in Fig 5A. As previously reported, the ability to utilize arabinotriose was abolished in the Δ*msmX* mutant, as evidenced by the drastic increase in doubling time (11). Conversely, the growth phenotype was rescued in both the ΔS1254 Δ*msmX* and *frlRstop* Δ*msmX* double mutants, even though the doubling time was slightly higher than the WT (Fig. 5A). Similarly, and as already reported (12, 25), ectopic expression of *frlP* under the synthetic promoter P_spank-hy_ leads to the restoration of the growth phenotype (Fig. 5A). To analyze transcription, the levels of *frlP* mRNA in these mutant cells were measured by quantitative reverse transcription polymerase chain reaction (RT-qPCR) in the presence of arabinose and the obtained data were used to calculate gene expression fold-change relative to the WT (Fig. 5B). The mRNA levels of *frlP* in the Δ*msmX* single mutant were like those of the WT, indicating that this mutation does not affect *frlP* mRNA accumulation. Correlating with the functionality assay there was an increase of *frlP* mRNA levels in the constructed double mutants, which indicates that FrlP may be functionally substituting MsmX in the uptake of arabinotriose. Moreover, there is a significant difference between the amount of *frlP* mRNA measured in each double mutant (ΔS1254 Δ*msmX* vs *frlRstop* Δ*msmX*, Fig. 5B). The deletion ΔS1254 led to a higher amount of *frlP* mRNA level than that measured in the *frlRstop* mutant. These results point to both the direct involvement of FrlR and the indirect contribution of S1254 in the regulation of *frlP* expression. Western blot analysis performed in protein extracts from these strains corroborated the results of the functional analysis and RT-qPCR as FrlP accumulates in the strains where the growth phenotype was restored and for which the *frlP* transcript level increased (Fig. 5C). Interestingly, although a much higher increase of mRNA accumulation was noticeable in the mutant that expresses *frlP in trans* (Fig. 5B), a sharp decrease in doubling time was not observed (Fig. 5A), nor did it lead to increased protein accumulation (Fig. 5C), which could be due to a high transcript turnover (discussed below). These data show that in natural conditions, FrlP can substitute MsmX as NBD of the AraNPQ importer when FrlR is downregulated.

**Fig. 5.**
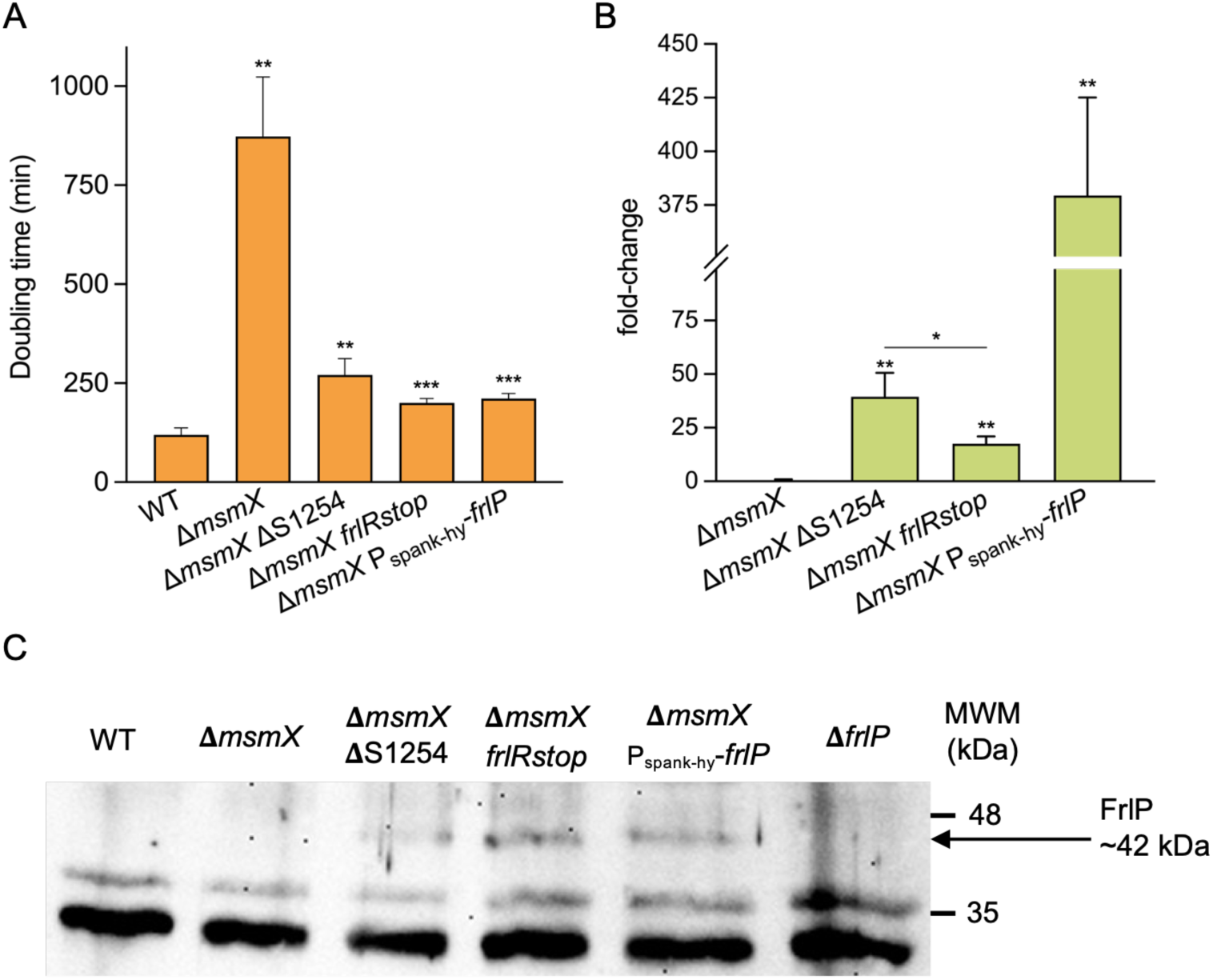
**(A)** Growth kinetic parameters of *B. subtilis* WT and mutants Δ*msmX* (IQB495), Δ*msmX* ΔS1254 (ISN51), Δ*msmX frlPstop* (ISN72) and *P_spank(hy)_ frlP* (ISN15) were measured in CSK minimal medium supplemented with 0.1 % (wt/vol) arabinotriose as sole carbon and energy source and are shown as doubling time (min). Values above 500 min were considered no growth (12, 25). The results represent at least three independent experiments and error bars represent standard deviation of the mean. **(B)** Representation of relative expression of *frlP* in the respective strains compared to the WT in CSK minimal medium supplemented with 0.1 % (wt/vol) arabinose. Primers ARA925 and ARA926 were used for *frlP* and fold-change was normalized using primers ARA583 and ARA584 for the 16S gene. Error bars represent standard deviation of the mean from C_t_ values of at least three independent assays. **(C)** Western blot analysis of FrlP (c.a. 41.4 kDa) accumulation in total cell extracts of different *B. subtilis* strains. *P*_spank(hy)_-*frlP* (ISN15) and Δ*frlP* (IQB618) were used as positive and negative controls, respectively. NZYColour Protein Marker II (NZYTech) was applied as standard, and it is partially represented (MWM). The uncropped image of this bolt is presented in Fig. S3. Statistical significance of doubling time and fold-change of different strains compared to the respective controls is indicated (*p<0.05; **p<0.01; ***p<0.001).

### FrlP is restricted to the *Bacillus subtilis* group

To elucidate the distribution of ABC type I NBDs FrlP and MsmX proteins amongst the *Bacillaceae* family, we retrieved all representative proteomes from species belonging to this family (629, as of 23 January 2023) (Table S1) and inferred a phylogenomic tree using 100 single-copy orthogroups. The resulting species tree (Fig. 6A and Fig. S4) recapitulates the phylogenetic relationship among the main lineages within *Bacillaceae* (35).

**Fig. 6.**
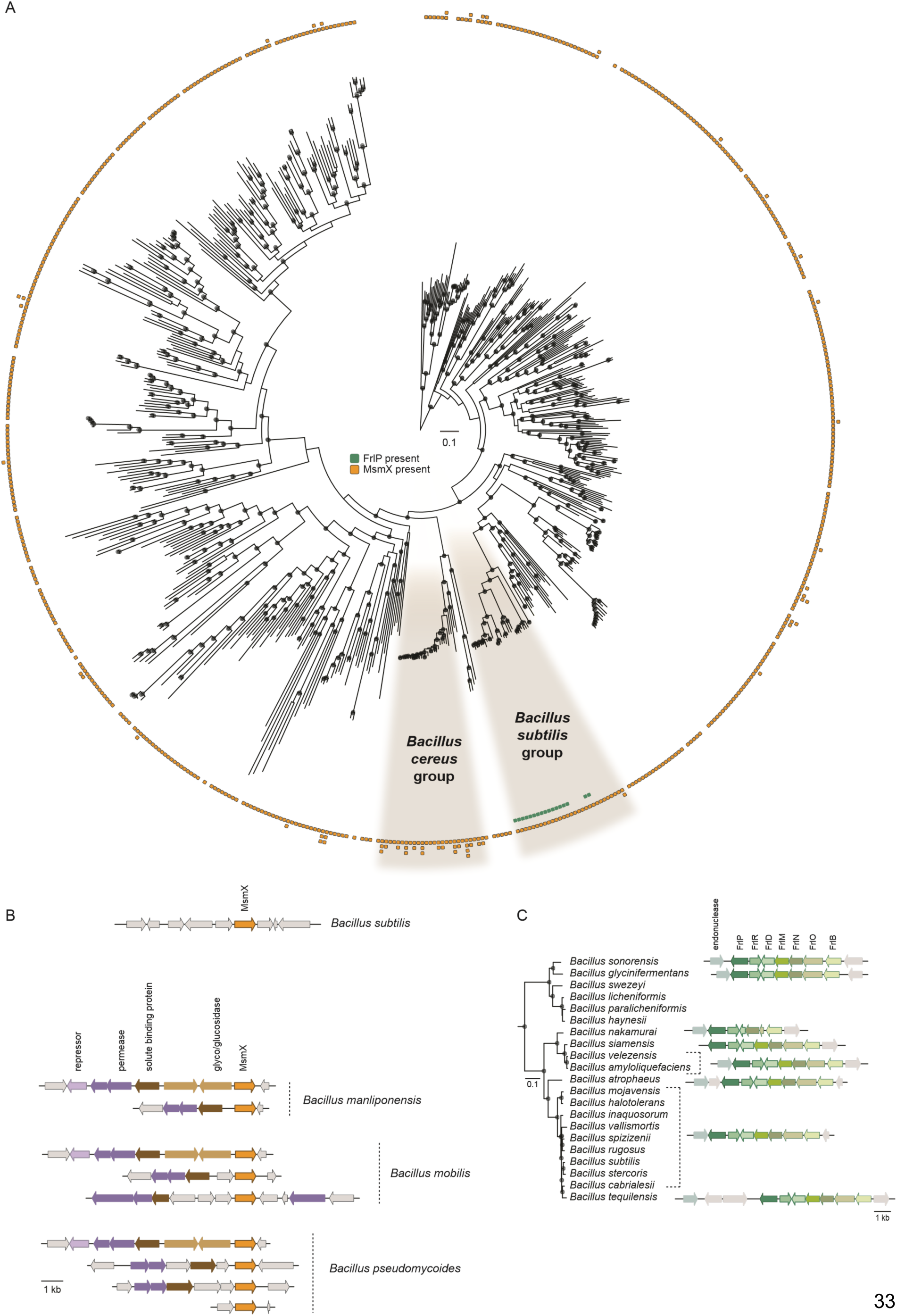
Phylogenomic tree of the *Bacillaceae*. **(A)** Maximum likelihood phylogenomic tree comprising 629 species from the *Bacillaceae* family inferred from the concatenated alignment of 100 single-copy orthogroups and rooted with *Staphylococcus aureus*. The species from the *Bacillus subtilis* and *Bacillus cereus* groups are highlighted. Presence/absence of *msmX*/*frlP* and number of protein-coding genes are depicted in the ring area. **(B)** Synteny of the *msmX* gene in *B. subtilis* and in three species of the *Bacillus cereus* group that contain more than one copy of the gene: *B manliponensis*, *B. mobilis* and *B. pseudomycoides*. Gene products are designated as annotated in the respective genomes. Genes sharing the same color encode proteins with similar predicted functions; grey arrows represent genes that are not related to sugar transport and, therefore, are not relevant for this study. **(C)** Pruned tree of the *Bacillus subtilis* group and respective genomic organization of the *frlP* cluster for each species. Orthologous genes are represented by the same color.

To further assess the distribution of the two ABC type I NBDs (FlrP and MsmX) among the *Bacillaceae* family we employed an HMM (hidden Markov model)-based strategy (36). Given the high similarity between the two NBDs (over 55%), a phylogeny containing all the recovered FlrP- and MsmX-like proteins was subsequently reconstructed to assess orthology (Fig. S5).

Of all the 629 different species, *Bacillus subtilis* and the *Bacillus cereus* groups stand out by displaying the most differences in the pattern of presence/absence of proteins. We noted that MsmX is virtually present in almost all species analyzed, although the *B. cereus* group is particularly enriched in MsmX paralogs (Fig. 6A), which may suggest an adaptive advantage for these species. The representation of *msmX* synteny in three different genomes of species belonging to the *B. cereus* group (Fig. 6B) shows a different genomic organization compared to that of *B. subtilis* 168, in which *msmX* is isolated in the genome as an orphan ATPase (7, 11). Contrarily, in *B. manliponensis*, *B. mobilis* and *B. pseudomycoides*, *msmX* is located next to or in the vicinity of genes that code for components of an ABC importer, with at least one conserved synteny amid all three species. On the other hand, FrlP is strictly present in the *B. subtilis* group, in 17 different species (Fig. 6A). The FrlP distribution in the phylogenetic tree suggests that an operon loss occurred in the most recent common ancestor of *B. swezey*, *B. licheniformis*, *B. paralicheniformis* and *B. haynesii*. The *frlBONMD*-*frlR*-*frlP* cluster synteny is conserved in all species, except in *B. nakamurai* in which gene loss and pseudogenization appears to be occurring (Fig. 6C). Furthermore, all organisms that contain FrlP, also contain MsmX, suggesting the importance of the latter to *B. subtilis* fitness.

The ecological association of species containing FrlP and MsmX duplications, excluding *B. subtilis sensu stricto*, was assessed by looking at the substrate of isolation of all genome entries for each species available on NCBI (BioSamples, between May and September 2024). Classification of environmental association was organized in five main groups: soils and plants, foods and beverages, host-associated, aquatic and other. Except for *B. glycinifermentans* and *B. sonorensis* that were mostly isolated in host-associated settings, organisms belonging to the *B. subtilis* group that contain FrlP were mainly isolated from soil and plant environments. As for organisms containing MsmX duplications, the main substrates of isolation were more widespread through other environments (Table S2). By looking at this simplified classification and at the results presented above we can infer that FrlP may be or may have been important for Amadori utilization, especially in plant-related surroundings.

## Discussion

In the present study, the physiological importance of the FrlP ABC-type I NBD (ATPase) was investigated. Information concerning FrlP has only been presumed through its genomic context. Hereby, by generating specific mutations, we show that the FrlMNO importer is required to utilize fructosevaline (Fig. 2). Moreover, not only FrlP is able to energize the FrlONM transporter, but so is MsmX, and both NBDs are functionally exchangeable in the presence of fructosevaline (Fig 2). These observations are supported by *in vivo* protein-protein interaction assays in *E. coli* confirming that both ATPases had affinity towards TMDs FrlN and FrlM, and that the interaction was stronger with the latter (Fig. 3). This unexpected finding extends the role of MsmX as major multitask ABC-type I ATPase responsible for carbohydrate import in *B. subtilis* (10–14). Previous reports have shown that *frlP* is expressed in a long transcript that include *frlBONMD-frlP* (30) driven by the promoter P*_frlB_* (16, 32, 33). However, two other shorter transcription units were also identified (17) (Fig. 1). Contradictory statements concerning the specific regulation of the amadori operon by the transcription factor FrlR in *B. subtilis* are found in the literature (16, 28). Thus, the construction of a nonfunctional FrlR mutant was imperative to clarify this subject and study the expression of *frlP*. Two lines of evidence indicate that this protein acts as a repressor: (i) the *frlR* nonsense mutation retards the *lag* phase of cells growing in fructosevaline relative to the WT (Fig. 2) and (ii) the same mutation increases expression of 5’*frlP*-*lacZ* fusions constructed in its own locus when compared to the WT (Fig. 4). The extended *lag* phase in the WT might be due to a presumably delay in Amadori metabolization, as observed for the FrlR homolog in *E. coli* (34). In the absence of fructosevaline the mutant was highly derepressed both at transcriptional and translational levels compared to the WT. The availability of fructosamines most likely decreases the affinity of FrlR to the DNA leading to the derepression of the operon and the consequent expression of *frlP*. The mechanism by which FrlR from *E. coli* is derepressed was proposed to be like that of NagR (YvoA) from *B. subtilis*, a GntR type transcription factor that negatively regulates genes from the N-acetylglucosamine (GlcNAc)-degrading pathway (34, 37). Phosphorylated effectors GlcNAcP or glucosamine-6-phosphate (GlcNP) allosterically bind to the C-terminal region of NagR, weakening the DNA binding ability of the winged helix-turn-helix (wHTH) DNA binding domain in the N-terminal region and, therefore, abolishing the repression that was being exerted (37, 38). Graf von Armansperg and colleagues proposed that fructoselysine-6P is the cognate substrate of FrlR in *E. coli* (34). Given the high similarity between the FrlR from *B. subtilis* and the homolog in *E. coli*, including the conservation of some phosphate binding residues (34), FrlR from *B. subtilis* may recognize phosphorylated Amadori compounds as well. This would cause the abolishment of its DNA binding ability and thus allowing the expression of downstream genes necessary for Amadori catabolism, including *frlP*. Deppe *et al*. 2011 showed that Amadori composed by the condensation of glucose and arginine could support growth in *B. subtilis* (16). Here, we found that *B. subtilis* can grow in the presence of another type of Amadori product, fructosevaline, which corroborates the findings of Wiame et al. 2004 which analyzed the in vitro properties of the FrlB and FrlD enzymes of *B. subtilis* indicating their role in the metabolism of fructosamines linked to the α-amino group of amino acids (24) pointing to the capacity of metabolize a number of different substrates. Thus, we propose that the correct name for the *frlBONMD-frlP* operon in *B. subtilis* is fructosamines [or Amadori operon; (16)] instead of fructoselysine operon (28).

Several functional studies have established MsmX as a multitask NBD, since it is the energy generator component of several ABC-type I sugar importers (10–14). In this study we show that both MsmX and FrlP participate as NBD of the fructosamines importer FrlONM (Fig. 7). Moreover, our laboratory showed that FrlP can mechanistically replace MsmX function in the transport of arabinotriose (AraNPQ) and galactan (GanSPQ) when ectopically expressed (12, 25) and pointed to the existence of a genomic context-dependent posttranscriptional mechanism involved in the regulation of *frlP* expression (12). To address this question, we target a putative antisense RNA S1254 located in transcriptional unit *frlBONMD*-*frlP* positioned between genes *frlD* and *frlP* and overlapping the entire regulator gene *frlR*, which is transcribed in the opposite direction (Fig. 1). The analyses of strains carrying mutations ΔS1254 or *frlRstop* by determination of growth kinetics parameters, RT-qPCR and western blot (Fig. 5) show that accumulation of FrlP is under the dual control of FrlR and S1254 RNA. Absence of FrlR (frlRstop mutation) when compared to both FrlR and S1254 RNA (ΔS1254 mutation) indicates that in addition to the specific direct control of *frlP* transcription by FrlR, the S1254 RNA plays a direct or indirect contribution in FrlP expression (Fig. 5). It is plausible to hypothesize a role of the antisense S1254 RNA in the autoregulation of the repressor FrlR by sequestering *frlR* mRNA and consequently affecting *frlP* expression. Alternatively, or in addition, it may provide a double stranded RNA upstream from *frlP* for RNase III-mediated digestion of overlapping transcripts, offering a mechanism to adjust mRNA levels of *frlP*. The latter posttranscriptional mechanism observed in several Gram-positive bacteria (39), will contribute to the maintenance of appropriate levels of FrlP in response to nutrient availability. Thus, our findings add another level of regulation to the complex posttranscriptional regulation of the fructosamines operon in *B. subtilis*. Besides the specific regulation by the FrlR repressor in response to fructosamines and the repression of the general transcription factor CodY triggered by branched-chain amino acids, both exerted at the transcriptional level, other factors, such as YlxR (nucleoid-associated protein modulator of RNA polymerase), RNaseY (essential 5’-end sensitive endoribonuclease) and CshA (*B. subtilis* major RNA helicase) are known to directly or indirectly participate in the regulation of the *frlBONMD-frlP* operon (15, 28, 32, 33, 40). The fructosamines operon is subject to the severe YlxR-dependent transcription repression (41). At the mRNA level, RNaseY likely degrades mRNA of the *frlBONMD-frlP* operon since its depletion leads to increase of mRNA of operon (40) and depletion of CshA strongly affects *frlBONMD-frlP* expression leading to mRNAs decrease (30). Similarly to that postulated by Deppe and colleagues, 2011 for the fructosamines operon (16), and based on the results obtained in this study (Fig. 4), we propose that during growth on glucose (or fructose), intracellular glucose 6-phosphate leads to the glycation of proteins and amino acids, fructosamine 6-phosphates inactivates FrlR, which consequently increases expression of FrlP. During growth on casaminoacids expression of FrlP is decreased via CodY-dependent transcription repression. Conversely, during growth on glucose expression of *msmX* is subjected to repression under the control of global transcriptional repressor CcpA (42). During growth on casaminoacids *msmX* expression is increased (43). This ensures that a functional NBD MsmX or FrlP is present in the cell to energize the fructosamines importer FrlONM (Fig. 7). Since FrlP is also able to substitute MsmX as energy motor for the AraNPQ and GanSPQ it would be interesting to investigate if the same is valid to the remaining ABC type-I importers energized by MsmX (Fig. 7). Remarkably, MsmX and FrlP display a structural C-terminal domain fold like the one present in MalK from *E. coli* that recruits regulatory proteins (27), suggesting additional levels of regulation targeting the two NBDs. Noteworthy, both NBDs FrlP and MsmX are encoded in operons belonging to a class of metabolic operons that are heterogeneously expressed across a clonal population of isogenic siblings (28, 44). Bacterial populations engage in bet-hedging strategies to maximize survival since heterogeneity increases fitness when compared with homogeneous populations (45, 46). The differential expression of FrlP and MsmX in subsets of cells may allow the concomitant import of different carbon sources by the total population (Fig. 7).

**Fig. 7.**
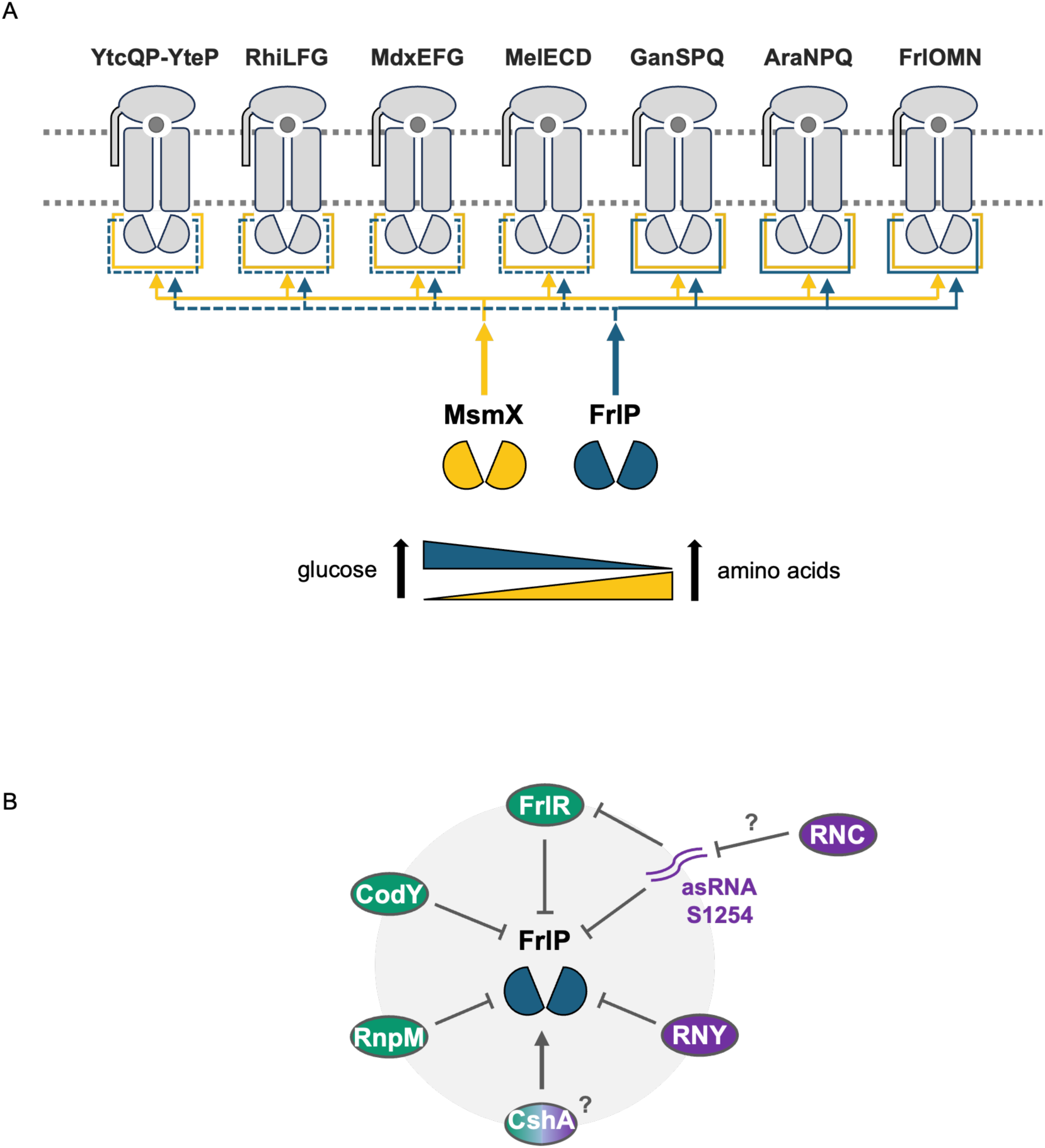
Model for the regulation of FrlP and its impact in the physiology of B. subtilis. **(A)** Interplay between NBDs MsmX and FrlP and sugar ABC-type I importers in *B. subtilis*. Full lines represent experimentally verified functional energization of the respective transporters; dashed lines represent putative binding of the NBD to the respective transporters. Below is depicted the intracellular concentration of MsmX and FrlP is response to nutrient availability regulated by global transcription factors CcpA and CodY. In the presence of glucose (left) as carbon and energy source intracellular accumulation of MsmX (depicted by a yellow triangle showing increasing concentration from left to right) is low due catabolite repression of *msmX* exerted by CcpA at the transcriptional level. On the other hand, in response to amino acids (right) as source of carbon and energy the level of FrlP (depicted by a blue triangle pointing decreasing concentration from left to right) is low due to the action of CodY which transcriptionally downregulates *frlP* expression. **(B)** Regulators involved in *frlP* expression at transcriptional (green) and posttranscriptional (purple) levels. FrlR and antisense asRNA S1254 both contribute to the decrease of *frlP* expression and consequent protein accumulation. We hypothesize that the asRNA S1254 regulates the operon repressor FrlR by sequestering *frlR* mRNA, furthermore, the overlapping transcripts S1254/*frlR* may provide a target for mRNA degradation and adjustment by RNaseIII (RNC). The global transcriptional regulator CodY binds and represses the *frl* operon expression in the presence of amino acids (16, 32, 33). RNaseY (RNY), which makes part of the putative RNA degradosome, breaks down the mRNA of the *frlBONMD*-*frlP*(*yurJ*) operon (40). RNA helicase CshA depletion leads to increased levels of *frlBONMD*-*frlP* transcripts (30) suggesting indirect transcriptional effects rather than RNA decay-related activity. Conversely, RnpM (formerly YlxR) strongly represses *frlBONMD*-*frlP* expression at the transcriptional level (28).

*Bacillus* spp, including *Bacillus subtilis*, represent the major genus of the Gram-positive bacterial populations in the soil and rhizosphere, and one of the most widely spread endophytic bacteria (47). Additionally, evidence indicates that *B. subtilis* is part of the normal gut microbiota of animals including humans (48, 49). Although research towards understanding Amadori utilization in bacteria is scarce, the catabolic machinery is complex and involves a different set of enzymes and is present in several organisms found in soils, rhizosphere, and associated to the gastrointestinal tract (19, 50–55). Examples include the capacity to utilize amadori deoxyfructosyl glutamine, which is dispersed through bacteria of the family *Rhizobiaceae* commonly found in rhizospheres and rotting plant materials (19, 50) or fructoselysine degraded by human intestinal microbiota (51–53). To assess the importance of the presence of FrlP in the cell, given the promiscuous nature of MsmX binding, the phylogenetic distribution of both FrlP and MsmX amongst the *Bacillaceae* family was investigated. FrlP seems to be restricted to the *B. subtilis* group, while MsmX is phylogenetically well distributed in the *Bacillaceae* family tree, and MsmX duplication events seem to be enriched on the *B. cereus* group (Fig. 6A). Regarding our data about the ecological classification of the *B. subtilis* group and other *Bacillaceae* family member that contain duplications of MsmX, it should be noticed that the isolation source of a given organism may not correspond to the preferred environments of a given species; nevertheless, it can give us hints regarding possible ecological niches (56). The presence of FrlP and the respective Frl components of the fructosamines operon in the *B. subtilis* group, together with the identification of other soil organisms that can degrade fructosamines, suggests that this trait is conserved among organisms that are found associated to plants and soil, highlighting their nutritional relevance and contribution to bacterial fitness.

## Materials and Methods

### DNA Manipulation and Sequencing

Regular DNA manipulations were performed as previously described by Sambrook *et al.* (57). PCR amplifications were carried out using Phusion High-Fidelity DNA Polymerase from Thermo Fisher Scientific according to the manufacturer’s recommendations. Restriction enzymes, FastAP Thermosensitive Alkaline Phosphatase and T4 DNA ligase used for restriction cloning purposes were purchased from Thermo Fisher Scientific and were also used according to the manufacturer’s instructions. DNAs were purified from agarose gel bands or from PCR and restriction reaction mixtures with the NZYGelpure kit (NZYtech). Plasmid DNA was extracted and purified using the NZYMiniprep kit from NZYtech. Plasmids and PCR products were verified by restriction patterns and/or Sanger sequencing at STAB VIDA (https://www.stabvida.com/). All primers and plasmid DNA used in this work are listed on Table S3 and Table S4.

### Construction of *B. subtilis* strains

The generation of the markerless ΔS1254 and *in-frame* Δ*frlO* deletion mutants, together with the *frlRstop* mutant in the chromosome of *B. subtilis* was achieved by overlap extension PCR followed by allelic replacement using a pMAD vector as described by Arnaud *et al*. (58). In brief, the immediately upstream and downstream regions of the target DNA were cloned together in the pMAD vector which was integrated in the chromosome via a single recombination event, promoted by growth at a non-permissive temperature for plasmid replication. The removal of the plasmid was upheld by a second recombination event that occurred in permissive temperature without antibiotic, allowing the withdrawal and the curing of the plasmid and the substitution of the targeted allele. The allelic replacements generated an 884 bp deletion of the S1254 region and an in-frame Δ*frlO* deletion of 525 bp that comprises amino acid-coding positions 121 to 295. The creation of the *frlRstop* mutation (E36stop) was obtained by overlap PCR using mutagenic primers (Supplemental data and Table S1), and the allelic replacement was done using the pMAD vector as described above. The *frlRstop* mutant resulted from the exchange of codon GAA (glutamic acid) to TAA (stop codon) and was confirmed by Sanger sequencing. The ΔS1254 deletion was made in a Δ*msmX*::*cat B. subtilis* background, and the *frlRstop* mutation was introduced in WT and Δ*msmX*::*cat* strains. Transcriptional and translational fusions of the gene *lacZ* of *E. coli* to the 5’ region of *frlP* were created by a single recombinational event at the respective locus of *B. subtilis* chromosome using integrative plasmids that contained about 340 bp of the upstream region of *frlP* plus the respective 5’*frlP*-*lacZ* fusions, with and without the *frlRstop* mutation. Transformation of the constructed *B. subtilis* strains was performed according to the methodology described by Anagnostopoulos and Spizizen (59). All *B. subtilis* strains used and constructed in this work are listed in Table 1. A detailed description of the construction of *B. subtilis* strains is found in the Supplementary Data section; primers and plasmids are listed in Table S3 and Table S4, respectively.

**Table 1.** List of strains used and constructed for this work.

| Strain | Relevant Genome | Source |
| --- | --- | --- |
| ***E. coli*** |  |  |
| XL10 Gold | Tet<sup>r</sup>Δ (*mcrA*)183 Δ (*mcrCB-hsdSMR-mrr*)173 *endA1 supE44 thi-1 recA1 gyrA96 relA1 lac* Hte [F' *proAB lacI<sup>q</sup>ZΔM15* Tn10 (Tet<sup>r</sup>) Amy Cam<sup>r</sup>] | Stratagene |
| BL21(DE3)pLysS | F- *ompT hsdS<sub>B</sub>(r<sub>B</sub><sup>-</sup> m<sub>B</sub><sup>-</sup>) gal dcm* (DE3) pLysS (Cm<sup>R</sup>) | (68) |
| BTH101 | F-, *cya-99, araD139, galE15, galK16, rpsL1 (Str<sup>r</sup>), hsdR2, mcrA1, mcrB1* | Euromedex |
| ***B. subtilis*** |  |  |
| 168 T<sup>+</sup> | Prototroph | F. E. Young |
| IQB495 | Δ*msmX::cat* | (11) |
| IQB618 | Δ*frlP(yurJ)* | (12) |
| IQB634 | Δ*frlP(yurJ)* Δ*msmX::cat* | (69) |
| ISN51 | Δ*msmX::cat* ΔS1254 | pIG1→IQB495 |
| ISN71 | *frlRstop36* | pIG6→168T<sup>+</sup> |
| ISN72 | *frlRstop36* Δ*msmX::cat* | pIG6→IQB495 |
| ISN88 | Δ*frlO* | pIT1→168T<sup>+</sup> |
| ISN128 | Φ(*frlP'-lacZ*+) (transcriptional fusion) *km* | pIG25→168T<sup>+</sup> |
| ISN129 | Φ(*frlP'-lacZ*+) (translational fusion) *km* | pIG26→168T<sup>+</sup> |
| ISN130 | *frlRstop36* Φ(*frlP'-lacZ*+) (transcriptional fusion) *km* | pIG27→ISN71 |
| ISN131 | *frlRstop36* Φ(*frlP'-lacZ*+) (translational fusion) *km* | pIG28→ISN71 |

### Growth conditions

*Escherichia coli* XL10-Gold (Stratagene), used for the construction of all plasmids, *E. coli* BL21(DE3)pLysS, used for heterologous expression of His_6_-FrlP (described in Supplemental Data), and *E. coli* BTH101, used for B2H assays, were all grown in liquid Lysogeny Broth (LB) medium (60) and on LB solidified with 1.6 % (wt/vol) agar. LB media were supplemented with ampicillin (100 μg mL^-1^), kanamycin (30 μg mL^-1^), tetracycline (12 μg mL^-1^), IPTG (0.1 or 0.5 mM) or streptomycin (100 μg mL^-1^) when appropriate.

*B. subtilis* strains were routinely grown in liquid LB medium, LB solidified with 1.6 % (wt/vol) agar or liquid SP medium (61), supplemented with chloramphenicol (5 μg mL^-1^), kanamycin (10 μg mL^-1^), erythromycin (1 μg mL^-1^) and/or X-Gal (80 μg mL-1), when suitable. Growth kinetic parameters of *B. subtilis* were determined in M9 minimal medium (60) without trace elements, supplemented with 1 mM D-(+)-glucose (Sigma), 1 mM D(-) fructose (Merck) and 2 mM fructosevaline (mixture of diastereomers) (CymitQuimica). Starter cultures were grown overnight in 5 mL M9 minimal medium supplemented with 27.8 mM glucose and 0.05 % (wt/vol) Casein acid hydrolysate, from bovine milk (CH) (Fluka) and diluted to an initial OD_600nm_ of 0.01 in 200 μL M9 minimal medium supplemented with 22 μg mL^-1^ CAF and with the respective carbon and nitrogen source on a 96 well plate. Growth was followed by measuring the optical density (OD_600nm_) in 15 min intervals for 12 h or 48 h (depending on the carbon source) in a microplate reader (Spectramax) at 37 °C with agitation. Growth curves were analyzed after normalizing the OD_600nm_ with blank absorbance values and were used to determine the doubling time of each strain, when growth phenotype was observed. Doubling time was calculated with absorbance values taken between 1 h and 3 h before each growth peak. Growth kinetic parameters of *B. subtilis* constructed strains were also determined in CSK minimal medium supplemented with 0.1 % (wt/vol) α-1,5-arabinotriose (Megazyme) as sole carbon and energy source, as described by Ferreira and Sá-Nogueira (11). Supplementation of CSK medium with IPTG (1 mM) was done when applicable.

### Bacterial Adenylate Cyclase Two-Hybrid (B2H) System

The Bacterial Adenylate Cyclase Two-Hybrid System ((31); Euromedex) was used to test protein-protein interactions between Nucleotide Binding Domains (NBDs) MsmX and FrlP and Transmembrane Domains (TMDs) FrlN and FrlM. ATPases were fused to fragment T18 in the pUT18 vector, while TMDs were fused to fragment T25 in pKT25 or pKNT25 plasmids. Construction of plasmids for B2H is described in the Supplementary Data section. *E. coli* BTH101 was co-transformed with a combination of one pUT18 and one pKT25 or pKNT25 derivatives; the resulting co-transformants were selected in LA supplemented with the appropriate antibiotics. Co-transformants were inoculated in LB with the respective antibiotics and IPTG (0.5 mM), grown aerobically at 30 °C and protein-protein interactions were assayed for β-galactosidase activity following kit instructions.

### β-galactosidase assays of *B. subtilis* strains

*B. subtilis* strains holding transcriptional and translational fusions of the 5’ region of *frlP* to *lacZ* gene were grown in M9 minimal medium in a 96 well plate supplemented with CAF plus glucose, fructose, or fructosevaline, as described above, or 1 % (wt/vol) casein acid hydrolysate from bovine milk (CH) (Fluka). 200 μL of each triplicate (600 μL total) were recovered to the same microcentrifuge tube at the approximate growth peak of each strain and kept at -20 °C until the β-galactosidase assay was performed the next day. Cells were resuspended in 1 mL Z buffer (60) with 100 μg mL^-1^ lysozyme and left incubating at 37 °C for 5 min. Triton X-100 0.1 % (wt/vol) was added to each tube and the cells were vigorously vortexed for 10 sec and incubated on ice for 5 min (adapted from (62)). The tubes were incubated at 28 °C for 10 min in a dry bath before starting the assay. β-Galactosidase activity was measured and expressed in Miller units (M.U.) as previously described, using the substrate ONPG (60). *B. subtilis* 168T^+^ (WT) (without *lacZ* fusions) was used as control; M.U. range between 1.4 and 2.7 in glucose, fructose and CH and between 5.1 and 12 in fructosevaline.

### RNA extraction and RT-qPCR assays

For total RNA extraction, B. subtilis strains were grown in 10 mL CSK minimal medium supplemented with 0.1 % (wt/vol) L-(+)-arabinose (Sigma) and 1.4 mL of cell cultures were collected at OD_600nm_ of 0.7 - 0.8 as previously described (12). Total RNA extraction was performed with the Absolutely Total RNA Miniprep kit (Agilent Technologies) according with manufacturer’s instructions using RNase-free procedures; RNA integrity was analyzed in 1 % (wt/vol) agarose TBE 1x gel; total RNA was quantified using a NanodropTM 1000 Spectrophotometer (Thermo Fisher Scientific) and kept at -80 °C in ready-to-use 10 μL aliquots. DNA contamination of RNA samples was assessed by standard PCR using primers ARA916 and ARA917. RT-qPCR assays were performed using primers ARA583 and 584 (16S gene) (12) and primers ARA925 and ARA926 (*frlP*). Primer efficiency was assessed using the Rotor-Gene SYBR® Green PCR Kit (Qiagen) in a Rotor-Gene 6000 (Corbett) real-time cycler and RT-qPCR experiments were performed using the SensiFAST™ SYBR No-ROX One-Step Kit (Bioline), in a Rotor-Gene Q MDx (Qiagen) real-time cycler. RT-qPCR was performed according to the manufacturer’s instructions, using 40 ng total RNA and 0.1 μM of each primer, in a total volume of 12.5 μL. This comparative analysis was performed using Ct values from at least two independent assays and the statistical analysis was performed on Prism 10 using an unpaired t test.

### Protein extracts preparation and Western Blot

*B. subtilis* strains were grown in the same conditions as previously described for RT-qPCR assays and cells were collected at the same time point. As positive and negative controls for FrlP accumulation an ectopically overexpressing FrlP strain (ISN15) under the control of a synthetic promoter (P_spank-hy_) (25) and an *frlP* deletion mutant (IQB618) were used, respectively. The strain containing *frlP* in trans was grown with an additional supplementation of 1 mM IPTG (isopropyl β-D-1-thiogalactopyranoside) (25). Total cell extracts and total protein quantification were done as previously described (12). 20 μg of total protein from each extract were loaded in a 12.5 % SDS-PAGE and run at constant electrical current (80 mA) for 50 min using NZYColour Protein Marker II (NZYTech) as standard. Fractionated proteins were transferred to a nitrocellulose membrane (0.45 μm; Bio-Rad) and visualized with Ponceau Red. The membrane was blocked with 5 % (wt/vol) powdered milk solution in TBS-Tween 1x for 1h, washed and incubated overnight with anti-His_6_-FrlP 1:1,000 in TBS-Tween with 0.5 % (wt/vol) powdered milk, followed by washing and 30 min incubation with HRP-conjugated goat anti-rabbit IgG antibody 1:10,000 (Thermo Scientific, Pierce Antibody Products). After a final washing step, the membrane was incubated with SuperSignal West Pico PLUS (Thermo Fisher Scientific) and blots were imaged using the Invitrogen iBright FL 1500 Imaging System (Thermo Fisher Scientific).

### Phylogenomic reconstruction of the Bacillaceae

A total of 629 reference genomes and respective proteomes from the *Bacillaceae* family and the outgroup *Staphylococcus aureus* were retrieved from NCBI (accessed on 23 January 2023, Table S1). These proteomes were used as input for Orthofinder v2.5.4 (63) which inferred orthogroups based on all-versus-all sequence similarity searches using DIAMOND, followed by multiple sequence alignment with MAFFT and gene trees using IQ-TREE (64). A species phylogeny was reconstructed from 100 orthogroups that contained single-copy genes in at least 48.6 % of species. The resulting phylogeny was rooted using *Staphylococcus aureus*.

### HMM-based search of FrlP and MsmX across *Bacillaceae* and respective genomic context

To search for MsmX and FrlP across all species, an HMM profile was first constructed. For that the reviewed protein sequences of MsmX (P94360) and FrlP (O32151) from *Bacillus subtilis* were retrieved from UniProt and used as queries for a BLASTp search in NCBI refseq database. Hits with an e-value below 0.001 were retained, up to a maximum of 100 hits per protein. These sequences were then aligned using MAFFT v7.407 (65), and HMM profiles for each gene were generated using HMMER v3.3.3 (66) within Orthofisher (36). The resulting HMM profiles were used to assess presence/absence of each gene and respective copy number using Orthofisher v1.0.5 (36) with default parameters. Given the high homology between the two proteins (MsmX and FrlP) we further confirmed orthology of the identified hits by Orthofisher by constructing a phylogeny. For that, all the identified protein sequences by Orthofisher (both MsmX and FrlP) were aligned with MAFFT and used to construct a phylogenetic tree with IQ-TREE. Absence of FrlP was confirmed by BLASTp searches against the proteomes of the following species: *B. haynesii*, *B. paralicheniformis*, *B. licheniformis*, *B. swezeyi*, *B. australimaris*, *B. safensis*, *B. pumilus*, *B. zhangzhouensis*, *B. stratosphericus*, *B. gobiensis*, *B. capparidis*, *M. crassostreae*, *M. mangrovi*, *D. indicus*, *N. cucumis*, *S. horikoshii* and *G. kaustophilus*. The best hits were then used in a reciprocal BLASTp in NCBI against *B. subtilis* 168. *B. velezensis*, *B. sonorensis* and *B. amyloliquefaciens* were used as positive controls for the presence of these genes. Additionally, in order to confirm that absences were not originated by annotation issues, tBLASTx searches were performed for *B. haynesii*, *B. safensis*, *B. stratosphericus*, and *M. mangrovi* against the respective genome assemblies. These analyses confirmed the absence of FrlP in all analyzed species. The same procedure was conducted for *msmX*, for which tBLASTx searches were performed against the genome assemblies of 18 species identified as either lacking *msmX* or containing only a single copy. The results pointed to the presence of *msmX* in 15 species. Therefore, it is important to note that the methodology employed may result in an underestimation of *msmX* presence. For species in which the presence of FrlP was confirmed and for representative species containing multiple MsmX copies, the genomic context was further analyzed. For this, the complete proteome was predicted using AUGUSTUS v3.3.3 (67), with the complete gene model and *Staphylococcus aureus* as reference. From the AUGUSTUS output, coordinates for the genes of interest were extracted and used to generate genomic plots in R using the genoPlotR package.

## Acknowledgments

The authors would like to thank L. Jaime Mota for critically reviewing the manuscript. This work was supported by FCT - Fundação para a Ciência e a Tecnologia, I.P., by grant PTDC/BIA-MIC/30696/2017 to I.S-N, grant PTDC/BIA-EVL/0604/2021 to C.G., and fellowship UI/BD/151153/2021 to I.C.G.. Computational resources were supported by grant 2023.09581.CPCA.A1 to A.P.. This work was further financed by national funds from FCT - Fundação para a Ciência e a Tecnologia, I.P., in the scope of the project UIDP/04378/2020 and UIDB/04378/2020 of the Research Unit on Applied Molecular Biosciences - UCIBIO and the project LA/P/0140/2020 of the Associate Laboratory Institute for Health and Bioeconomy - i4HB.

