## Supplementary Data for "FrlP, an ABC type I importer component of *Bacillus subtilis*: regulation and impact in bacterial fitness"

**Table S 2** Ecological distribution of deposited genomes on NCBI (BioSamples) of *Bacillaceae* family members that contain FrlP or more than one copy of MsmX. First two rows represent number of copies of MsmX and FrlP in the respective organism. The niche that is more representative for each organism is highlighted in lighter grey.

* *Bacillus subtilis* sensus stricto BioSamples were not analyzed because is well established as a soil and gastrointestinal commensal organism. It is an easily dispersible and persistent organism and, therefore, several Biosample entries are available.

| MsmX | FrlP | Organism | Total entries | Soil and Plants | Foods and beverages | Host-associated | Aquatic | Other | Unidentified /missing (%) |
| --- | --- | --- | --- | --- | --- | --- | --- | --- | --- |
| FrlP presence | | | | | | | | | |
| 1 | 1 | *Bacillus amyloliquefaciens* | *996* | 318 | 65 | 67 | 11 | 49 | 48.80% |
|  |  |  |  | 31.93% | 6.53% | 6.73% | 1.10% | 4.92% |  |
| 1 | 1 | *Bacillus atrophaeus subsp. globigii* | 4 | 1 |  |  |  |  | 75.00% |
|  |  |  |  | 25.00% | 0.00% | 0.00% | 0.00% | 0.00% |  |
| 1 | 1 | *Bacillus cabrialesii* | 5 | 5 |  |  |  |  | 0.00% |
|  |  |  |  | 100.00% | 0.00% | 0.00% | 0.00% | 0.00% |  |
| 1 | 1 | *Bacillus glycinifermentans* | 33 | 6 | 3 | 17 |  |  | 21.21% |
|  |  |  |  | 18.18% | 9.09% | 51.52% | 0.00% | 0.00% |  |
| 1 | 1 | *Bacillus halotolerans* | 92 | 58 | 10 | 3 | 4 |  | 18.48% |
|  |  |  |  | 63.04% | 10.87% | 3.26% | 4.35% | 0.00% |  |
| 1 | 1 | *Bacillus inaquosorum* | 141 | 31 | 7 |  |  |  | 73.05% |
|  |  |  |  | 21.99% | 4.96% | 0.00% | 0.00% | 0.00% |  |
| 1 | 1 | *Bacillus mojavensis* | 52 | 32 |  | 2 | 2 |  | 30.77% |
|  |  |  |  | 61.54% | 0.00% | 3.85% | 3.85% | 0.00% |  |
| 1 | 1 | *Bacillus nakamurai* | 5 | 4 |  |  |  | 1 | 0.00% |
|  |  |  |  | 80.00% | 0.00% | 0.00% | 0.00% | 20.00% |  |
| 1 | 1 | *Bacillus rugosus* | 3 | 2 |  | 1 |  |  | 0.00% |
|  |  |  |  | 66.67% | 0.00% | 33.33% | 0.00% | 0.00% |  |
| 1 | 1 | *Bacillus siamensis* | 57 | 30 | 5 | 5 | 3 | 2 | 21.05% |
|  |  |  |  | 52.63% | 8.77% | 8.77% | 5.26% | 3.51% |  |
| 1 | 1 | *Bacillus sonorensis* | 71 | 12 | 8 | 34 |  | 2 | 21.13% |
|  |  |  |  | 16.90% | 11.27% | 47.89% | 0.00% | 2.82% |  |
| 1 | 1 | *Bacillus stercoris* | 22 | 8 | 2 | 4 |  | 5 | 13.64% |
|  |  |  |  | 36.36% | 9.09% | 18.18% | 0.00% | 22.73% |  |
| 1 | 1 | *Bacillus subtilis** | 9247 |  |  |  |  |  |  |
| 1 | 1 | *Bacillus subtilis subsp. spizizenii* | 305 | 48 |  | 2 | 4 | 91 | 52.46% |
|  |  |  |  | 15.74% | 0.00% | 0.66% | 1.31% | 29.84% |  |
| 1 | 1 | *Bacillus tequilensis* | 21 | 9 | 1 | 6 | 2 |  | 14.29% |
|  |  |  |  | 42.86% | 4.76% | 28.57% | 9.52% | 0.00% |  |
| 1 | 1 | *Bacillus vallismortis* | 77 | 12 | 2 |  | 1 | 51 | 14.29% |
|  |  |  |  | 15.58% | 2.60% | 0.00% | 1.30% | 66.23% |  |
| 1 | 1 | *Bacillus velezensis* | 1604 | 675 | 157 | 140 | 47 | 159 | 26.56% |
|  |  |  |  | 42.08% | 9.79% | 8.73% | 2.93% | 9.91% |  |
| MsmX duplications | | | | | | | | | |
| 2 | 0 | *Alkalihalobacillus shacheensis* | 1 | 1 |  |  |  |  | 0.00% |
|  |  |  |  | 100.00% | 0.00% | 0.00% | 0.00% | 0.00% |  |
| 3 | 0 | *Aquibacillus albus* | 1 |  |  |  |  |  | 100.00% |
|  |  |  |  | 0.00% | 0.00% | 0.00% | 0.00% | 0.00% |  |
| 2 | 0 | Bacillus albus | 43 | 12 |  | 6 | 13 | 10 | 4.65% |
|  |  |  |  | 27.91% | 0.00% | 13.95% | 30.23% | 23.26% |  |
| 2 | 0 | Bacillus alveayuensis | 4 |  |  |  | 3 |  | 25.00% |
|  |  |  |  | 0.00% | 0.00% | 0.00% | 75.00% | 0.00% |  |
| 2 | 0 | Bacillus capparidis | 3 | 1 |  |  |  |  | 66.67% |
|  |  |  |  | 33.33% | 0.00% | 0.00% | 0.00% | 0.00% |  |
| 2 | 0 | Bacillus clarus | 2 | 2 |  |  |  |  | 0.00% |
|  |  |  |  | 100.00% | 0.00% | 0.00% | 0.00% | 0.00% |  |
| 2 | 0 | Bacillus dafuensis | 3 | 1 |  | 1 |  | 1 | 0.00% |
|  |  |  |  | 33.33% | 0.00% | 33.33% | 0.00% | 33.33% |  |
| 2 | 0 | Bacillus gaemokensis | 2 |  |  |  | 2 |  | 0.00% |
|  |  |  |  | 0.00% | 0.00% | 0.00% | 100.00% | 0.00% |  |
| 3 | 0 | Bacillus luti | 11 | 2 |  | 5 | 3 |  | 9.09% |
|  |  |  |  | 18.18% | 0.00% | 45.45% | 27.27% | 0.00% |  |
| 2 | 0 | Bacillus manliponensis | 1 |  |  |  | 1 |  | 0.00% |
|  |  |  |  | 0.00% | 0.00% | 0.00% | 100.00% | 0.00% |  |
| 3 | 0 | Bacillus mobilis | 66 | 8 | 7 | 17 | 2 | 14 | 27.27% |
|  |  |  |  | 12.12% | 10.61% | 25.76% | 3.03% | 21.21% |  |
| 2 | 0 | Bacillus nitratireducens | 27 | 13 | 1 | 3 | 2 |  | 29.63% |
|  |  |  |  | 48.15% | 3.70% | 11.11% | 7.41% | 0.00% |  |
| 3 | 0 | Bacillus pacificus | 119 | 13 | 39 | 25 | 7 | 3 | 26.89% |
|  |  |  |  | 10.92% | 32.77% | 21.01% | 5.88% | 2.52% |  |
| 3 | 0 | Bacillus paramycoides | 76 | 15 |  | 1 | 2 | 38 | 26.32 |
|  |  |  |  | 19.74% | 0.00% | 1.32% | 2.63% | 50.00% |  |
| 2 | 0 | Bacillus paranthracis | 348 | 44 | 114 | 94 | 6 | 19 | 20.40% |
|  |  |  |  | 12.64% | 32.76% | 27.01% | 1.72% | 5.46% |  |
| 2 | 0 | Bacillus proteolyticus | 12 | 5 |  |  | 2 |  | 41.67% |
|  |  |  |  | 41.67% | 0.00% | 0.00% | 16.67% | 0.00% |  |
| 4 | 0 | Bacillus pseudomycoides | 147 | 125 | 1 | 4 |  |  | 11.56% |
|  |  |  |  | 85.03% | 0.68% | 2.72% | 0.00% | 0.00% |  |
| 2 | 0 | Bacillus rhizoplanae | 1 |  |  |  |  |  | 100.00% |
|  |  |  |  | 0.00% | 0.00% | 0.00% | 0.00% | 0.00% |  |
| 2 | 0 | Bacillus testis | 2 |  |  | 1 |  |  |  |
|  |  |  |  | 0.00% | 0.00% | 50.00% | 0.00% | 0.00% | 50.00% |
| 2 | 0 | Bacillus wiedmannii | 258 | 147 | 35 | 26 | 6 | 3 | 15.89% |
|  |  |  |  | 56.98% | 13.57% | 10.08% | 2.33% | 1.16% |  |
| 2 | 0 | Bacillus xiapuensis | 2 | 2 |  |  |  |  | 0.00% |
|  |  |  |  | 100.00% | 0.00% | 0.00% | 0.00% | 0.00% |  |
| 2 | 0 | Fictibacillus nanhaiensis | 8 | 1 |  | 2 | 3 | 1 | 12.50% |
|  |  |  |  | 12.50% | 0.00% | 25.00% | 37.50% | 12.50% |  |
| 2 | 0 | Lederbergia citri | 1 | 1 |  |  |  |  | 0.00% |
|  |  |  |  | 100.00% | 0.00% | 0.00% | 0.00% | 0.00% |  |
| 2 | 0 | Litchfieldia alkalitelluris | 2 | 2 |  |  |  |  | 0.00% |
|  |  |  |  | 100.00% | 0.00% | 0.00% | 0.00% | 0.00% |  |
| 2 | 0 | Lysinibacillus halotolerans | 2 | 1 |  |  |  |  | 50.00% |
|  |  |  |  | 50.00% | 0.00% | 0.00% | 0.00% | 0.00% |  |
| 2 | 0 | Lysinibacillus telephonicus | 2 |  | 1 |  |  | 1 | 0.00% |
|  |  |  |  | 0.00% | 50.00% | 0.00% | 0.00% | 50.00% |  |
| 2 | 0 | Lysinibacillus timonensis | 1 |  |  | 1 |  |  | 0.00% |
|  |  |  |  | 0.00% | 0.00% | 100.00% | 0.00% | 0.00% |  |
| 2 | 0 | Mesobacillus persicus | 1 |  |  |  |  |  | 100.00% |
|  |  |  |  | 0.00% | 0.00% | 0.00% | 0.00% | 0.00% |  |
| 2 | 0 | Oceanobacillus chungangensis | 1 |  |  |  | 1 |  | 0.00% |
|  |  |  |  | 0.00% | 0.00% | 0.00% | 100.00% | 0.00% |  |
| 2 | 0 | Oceanobacillus polygoni | 3 |  | 2 |  |  |  | 33.33% |
|  |  |  |  | 0.00% | 66.67% | 0.00% | 0.00% | 0.00% |  |
| 2 | 0 | Oceanobacillus rekensis | 1 | 1 |  |  |  |  | 0.00% |
|  |  |  |  | 100.00% | 0.00% | 0.00% | 0.00% | 0.00% |  |
| 2 | 0 | Oceanobacillus zhaokaii | 1 |  |  | 1 |  |  | 0.00% |
|  |  |  |  | 0.00% | 0.00% | 100.00% | 0.00% | 0.00% |  |
| 2 | 0 | Peribacillus simplex | 72 | 43 | 2 | 3 |  | 14 | 13.89% |
|  |  |  |  | 59.72% | 2.78% | 4.17% | 0.00% | 19.44% |  |
| 2 | 0 | Pontibacillus litoralis | 1 |  |  | 1 |  |  | 0.00% |
|  |  |  |  | 0.00% | 0.00% | 100.00% | 0.00% | 0.00% |  |
| 2 | 0 | Priestia aryabhattai | 156 | 81 | 3 | 8 | 6 | 32 | 16.67% |
|  |  |  |  | 51.92% | 1.92% | 5.13% | 3.85% | 20.51% |  |
| 3 | 0 | Priestia endophytica | 20 | 9 |  | 5 |  | 2 | 20.00% |
|  |  |  |  | 45.00% | 0.00% | 25.00% | 0.00% | 10.00% |  |
| 2 | 0 | Priestia filamentosa | 15 | 9 |  |  | 2 | 1 | 20.00% |
|  |  |  |  | 60.00% | 0.00% | 0.00% | 13.33% | 6.67% |  |
| 3 | 0 | Priestia megaterium | 966 | 639 | 8 | 52 | 21 | 75 | 17.70% |
|  |  |  |  | 66.15% | 0.83% | 5.38% | 2.17% | 7.76% |  |
| 2 | 0 | Psychrobacillus lasiicapitis | 2 |  |  | 1 |  |  | 50.00% |
|  |  |  |  | 0.00% | 0.00% | 50.00% | 0.00% | 0.00% |  |
| 2 | 0 | Salibacterium aidingense | 2 |  |  |  | 1 |  | 50.00% |
|  |  |  |  | 0.00% | 0.00% | 0.00% | 50.00% | 0.00% |  |
| 2 | 0 | Salibacterium salarium | 2 | 1 |  |  |  |  | 50.00% |
|  |  |  |  | 50.00% | 0.00% | 0.00% | 0.00% | 0.00% |  |
| 2 | 0 | Sediminibacillus dalangtanensis | 1 | 1 |  |  |  |  | 0.00% |
|  |  |  |  | 100.00% | 0.00% | 0.00% | 0.00% | 0.00% |  |
| 2 | 0 | Sutcliffiella halmapala | 1 | 1 |  |  |  |  | 0.00% |
|  |  |  |  | 100.00% | 0.00% | 0.00% | 0.00% | 0.00% |  |
| 3 | 0 | Terrilactibacillus laevilacticus | 3 | 3 |  |  |  |  | 0.00% |
|  |  |  |  | 100.00% | 0.00% | 0.00% | 0.00% | 0.00% |  |
| 3 | 0 | Terrilactibacillus tamarindi | 1 | 1 |  |  |  |  | 0.00% |
|  |  |  |  | 100.00% | 0.00% | 0.00% | 0.00% | 0.00% |  |

**Table S3** List of oligonucleotides used in this work; modified nucleotides for point mutations and restriction sites are underlined.

| Oligonucleotide | Sequence (5’ - 3’) |
| --- | --- |
| ARA583 | TCGCGGTTTCGCTGCCCTTT |
| ARA584 | AAGTCCCGCAACGCGCGCAA |
| ARA826 | CAAAAAGGATCTTCACCTAGATCC |
| ARA837 | CTGATGGCTAGCTTAACATTTGAACACG |
| ARA871 | ATTATTGATGGATCCGTGCCG |
| ARA872 | GTTTGATGAATTCAGCTGACG |
| ARA873 | CGCTTCTGAAGATCCTTTGGTGTCATTTTTGTAGTAGG |
| ARA874 | CCTACTACAAAAATGACACCAAAGGATCTTCAGAAGCG |
| ARA916 | GGAATTCGGTCTGGTCATGC |
| ARA917 | CGAGAATACTATAAGCAGTTTCCAGTGACGAAATAACGG |
| ARA918 | CCGTTATTTCGTCACTGGAAACTGCTTATAGTATTCTCG |
| ARA919 | CGGGATCCGCTGCATACGAGCG |
| ARA925 | CGCCGAAAGAACGTGATATT |
| ARA926 | TCGCCATCTTTCTGAGCTTT |
| ARA929 | GCTCGTTTTAGGTAGGCAGC |
| ARA930 | GCTGCCTACCTAAAACGAGC |
| ARA932 | GGGAATTGCTTCTAGACTCAATATAAG |
| ARA933 | GGTTTTTTTGCTCTAGACGGATTCTCAC |
| ARA943 | GTTGTTTCTAGAGACGAAATAACGGG |
| ARA944 | GGAGCACCTTTTCTAGATACACAGCC |
| ARA945 | GTTTCCTCCTTGGTACCCCTTTGACTC |
| ARA951 | TTAAGGGGGAGGATCCAATGTTGCGG |
| ARA960 | ATGGGCGGATAAGCTTTGGCCG |
| ARA1030 | AATCTAGAGTGAAGCTTCGCACC |
| ARA1031 | ATTCTAGATACTCCTCCTTTCCGG |
| ARA1033 | CCGGTACCCCTTTTACACTGCCG |
| ARA1050 | GATCTAGAGATGCTGCCTCAGAAG |
| ARA1051 | ATTTCGCGAACGGGCAGACATGGC |
| ARA1052 | CAGAATTCCTGACGGTGATGCG |
| ARA1053 | AACCCGGGTCGACGTGTTCAAATGTTAATGAAGCC |

**Table S 4** List of plasmids used in this work.

| Plasmid | Relevant Construction | Source or reference |
| --- | --- | --- |
| pET28b(+) | Bacterial vector for the expression of N-terminal 6xHis-tagged proteins with a thrombin site, *kan* | Novagen |
| PJL3 | pLitmus29 derivative used as a source of kanamycin resistance cassette, *bla*, *kan* | (3) |
| pJM783 | Integrative vector containing *lacZ*, *bla*, *cat* | (2) |
| pKNT25 | B2H expression vector for N-terminal fusions to T25 fragment of CyaA, *kan* | (5) |
| pKT25 | B2H expression vector for C-terminal fusions to T25 fragment of CyaA, *kan* | (5) |
| pKT25-zip | pUKT25 derivative with zip fused to T25 fragment used as positive control for B2H, *bla* | (5) |
| pMAD | Plasmid used for allelic replacement in Gram-positive bacteria, *bla*, *erm* | (1) |
| pUT18 | B2H expression vector for N-terminal fusions to T18 fragment of CyaA, *bla* | (5) |
| pUT18-zip | pUT18 derivative with zip fused to T18 fragment used as positive control for B2H, *bla* | (5) |
| pIT1 | pMAD derivative used for Δ*frlO*, *bla*, *erm* | This work |
| pIG1 | pMAD derivative used for ΔS1254, *bla*, *erm* | This work |
| pIG20 | pKNT25 derivative for the expression of FrlN-T25 fusion protein, *kan* | This work |
| pIG23 | pKT25 derivative for the expression of T25-FrlM fusion protein, *kan* | This work |
| pIG24 | Integrative plasmid with a *lacZ* gene for the construction of transcriptional and translational fusions, *kan* | This work |
| pIG25 | Integrative plasmid used for the construction of *in situ* 5’*frlP*-*lacZ* (Φ(*frlP*’*-lacZ*+)) transcriptional fusion, *kan* | This work |
| pIG26 | Integrative plasmid used for the construction of *in situ* 5’*frlP*-*lacZ* (Φ(*frlP*’*-lacZ*+))translational fusion, *kan* | This work |
| pIG27 | Integrative plasmid used for the construction of *in situ* *frlRstop* 5’*frlP*-*lacZ* (Φ(*frlP*’*-lacZ*+)) transcriptional fusion, *kan* | This work |
| pIG28 | Integrative plasmid used for the construction of *in situ frlRstop* 5’*frlP*-*lacZ* (Φ(*frlP*’*-lacZ*+)) translational fusion, *kan* | This work |
| pIG6 | pMAD derivative used for the introduction of the nonsense mutation in *frlR* GAA (Glu at position 36) to TAA (stop codon), *bla*, *erm* | This work |
| pIG9 | pET28b(+)-based vector for the expression of His_6_-FrlP, *kan* | This work |
| pJS1 | pUT18 derivative for the expression of FrlP-T18 fusion protein, *bla* | This work |
| pLG55 | pUT18 derivative for the expression of MsmX-T18 fusion protein, *bla* | This work |
| pLG64 | pKT25 derivative for the expression of T25-AraQ fusion protein, *kan* | This work |

**Supplementary figures**

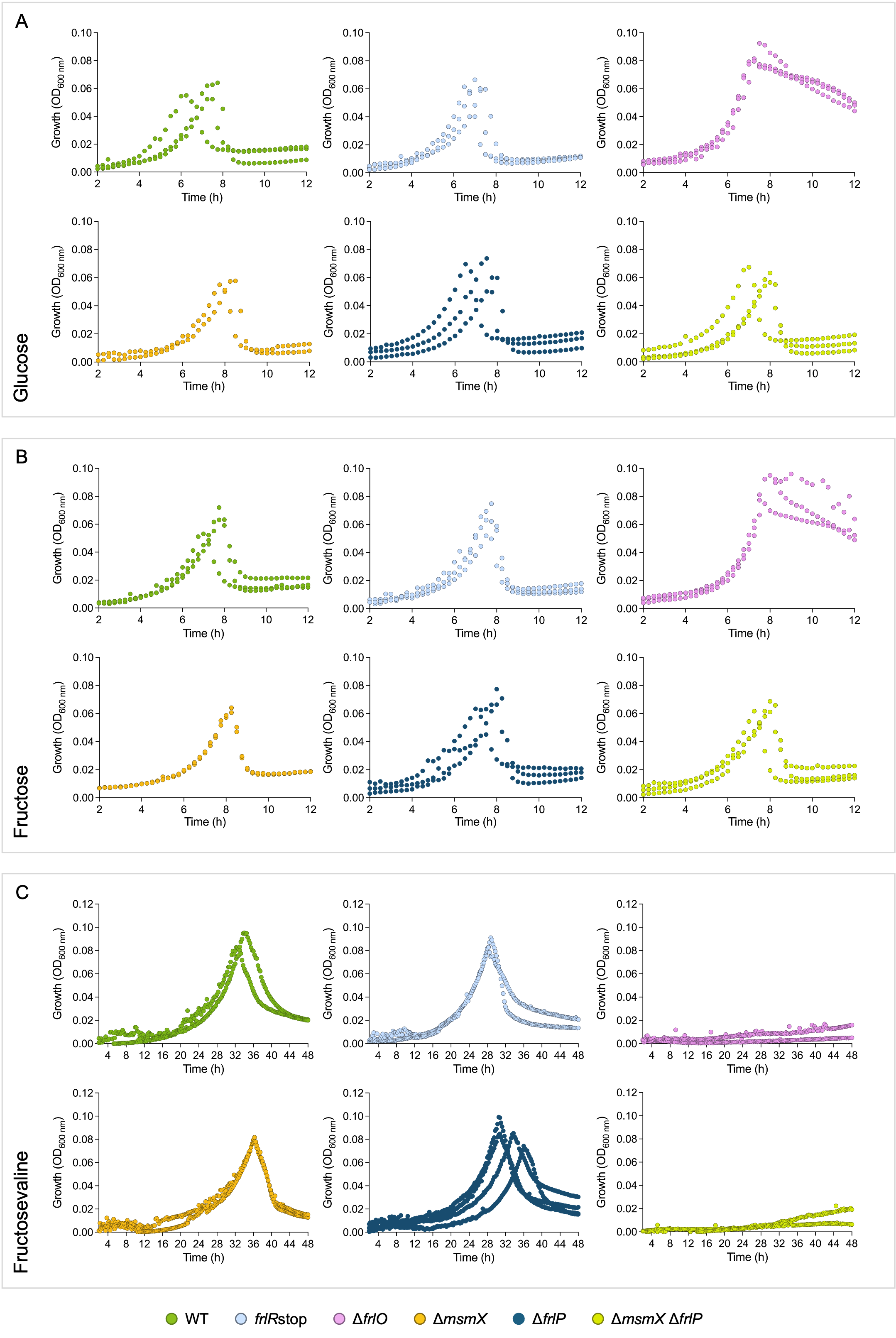

**Fig. S1.** Growth analysis of *B. subtilis* WT (darker green), *frlRstop* (light blue), Δ*frlO* (pink), Δ*msmX* (orange), Δ*frlP* (dark blue) and Δ*msmX*Δ*frlP* (light green) strains in M9 minimal medium supplemented with 22 μg mL^-1^ CAF and 1 mM glucose **(A)**, 1 mM fructose **(B)** or 2 mM fructosevaline **(C)**. At least two biological replicates are shown, each resulting from the mean of three technical replicates.

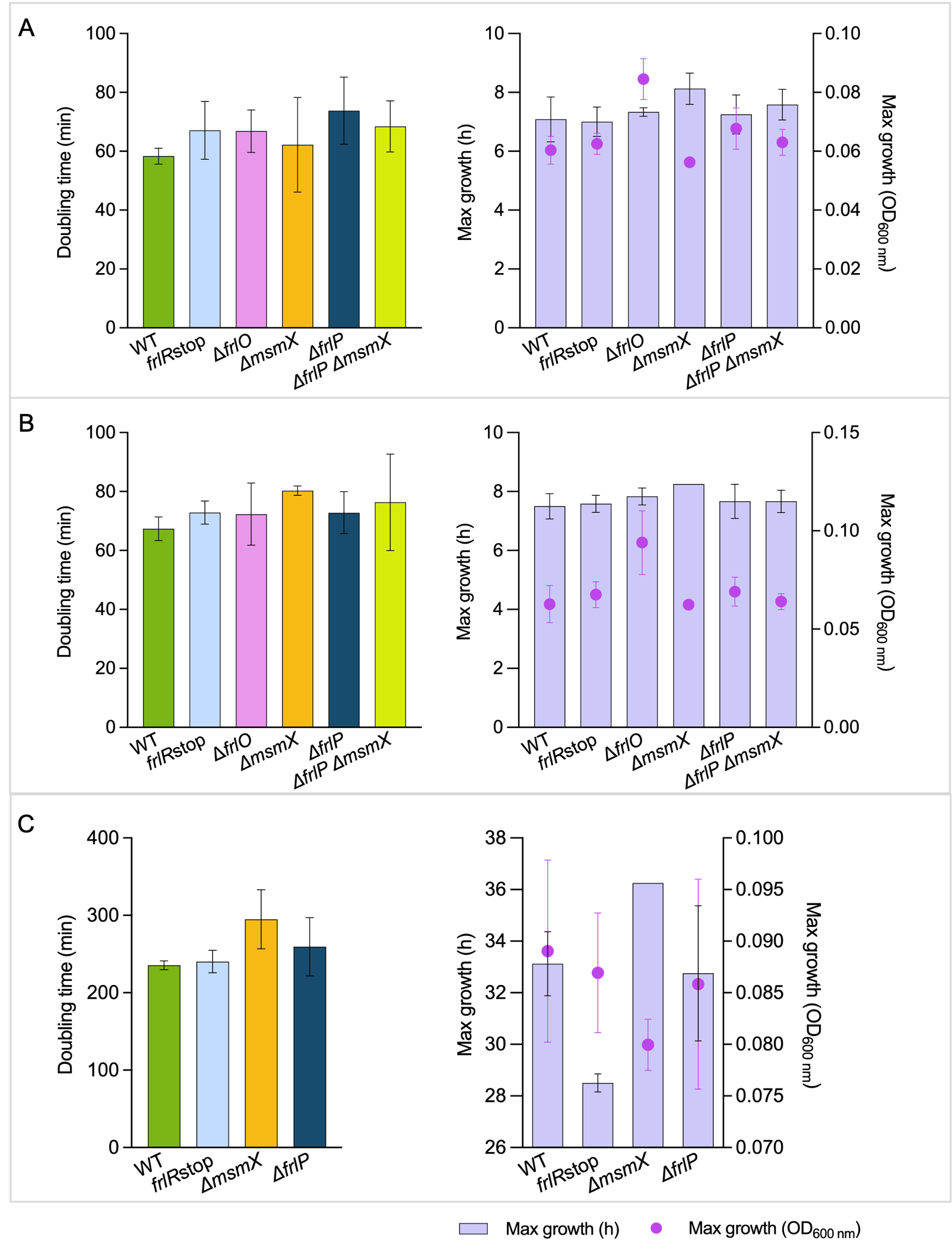

**Fig. S2.** Growth kinetic parameters of *B. subtilis* strains grown in M9 minimal medium supplemented with 22 μg mL^-1^ CAF and 1 mM glucose **(A)**, 1 mM fructose **(B)** or 2 mM fructosevaline **(C)**, from growth analysis shown in Figure S1. Left panels represent doubling time in minutes. Right panels represent maximum growth values, in hours of growth (bars) and in maximum OD_600nm_ values obtained (dots). Error bars represent standard deviation of the mean from at least two independent growth analyses.

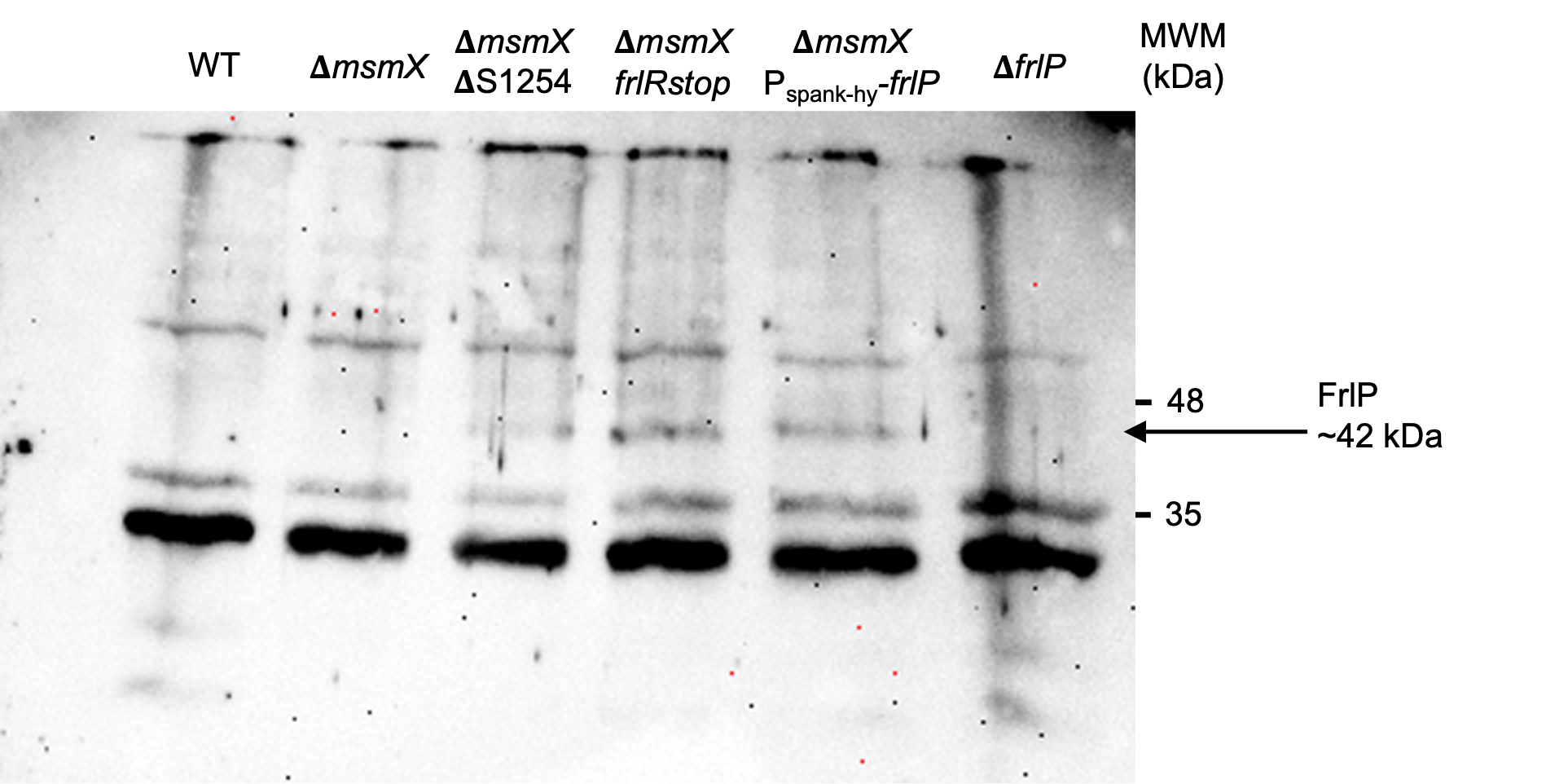

**Fig. S3.** Uncropped Western blot from Fig. 5C. Western blot analysis of FrlP accumulation in total cell extracts of different *B. subtilis* strains. *B. subtilis* Δ*msmX* P_spank-hy_-*frlP* and Δ*frlP* strains were used as positive and negative controls for FrlP accumulation, respectively. NZYColour Protein Marker II (NZYTech) was used, and it is depicted on the right.

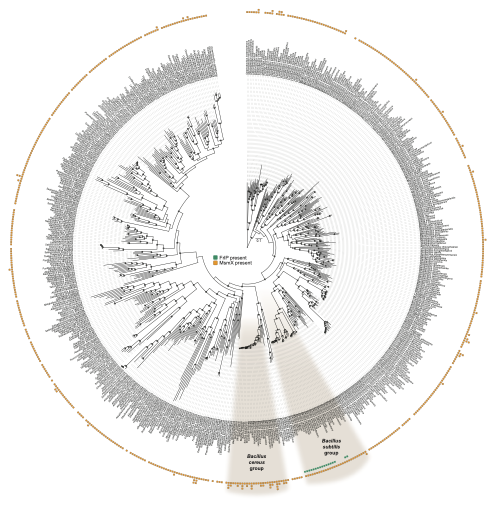

**Fig. S4.** Maximum likelihood phylogenomic tree comprising 629 species from the *Bacillaceae* family inferred from the concatenated alignment of 100 single-copy orthogroups and rooted with *Staphylococcus aureus*. The species from the *Bacillus subtilis* and *Bacillus cereus* groups are highlighted. Presence/absence of *msmX*/*frlP* and number of protein-coding genes are depicted in the ring area.

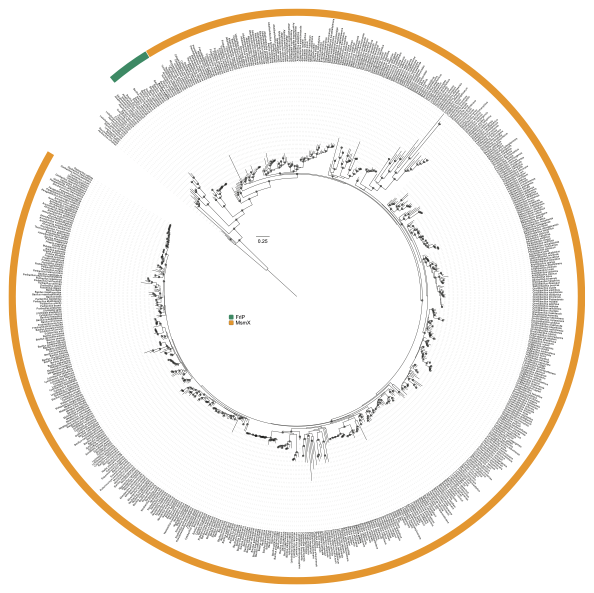

**Fig. S5.** Maximum likelihood phylogeny comprising the protein sequences retrieved from Orthofisher of MsmX and FrlP, depicted in orange and green, respectively. The reference sequences of FrlP and MsmX from *B. subtilis* are highlighted in **bold**.

### Construction of plasmids and *Bacillus subtilis* strains

The construction of markerless Δ*frlO* and ΔS1254, as well as the *frlRstop* mutation in the *B. subtilis* chromosome was obtained by allelic replacement using the pMAD vector preceded by overlap PCR, as described by Arnaud *et al.* (1). For deletion purposes primers were designed to amplify the upstream and downstream regions of the section to be deleted; mutagenic primers were used to create the nonsense point mutation in the *frlR* gene. For the construction of the Δ*frlO* mutant, chromosomal DNA of *B. subtilis* 168T^+^ was used for amplification of two PCR products with primers ARA871 and ARA873, and ARA872 and ARA874. The products from both PCR reactions were joined by overlap PCR using external primers ARA871 and ARA872; the resulting amplicon was digested with EcoRI and BamHI and cloned into pMAD digested with the same enzymes and dephosphorylated, yielding pIT1. ΔS1254 and *frlRstop* constructs were both made by overlap extension PCR using external primers ARA916 and ARA919. The ΔS1254 deletion and the *frlRstop* mutation were obtained using internal primers ARA917 and ARA918, and mutagenic primers ARA929 and ARA930, respectively. The mutagenic primers introduced the E36stop mutation in *frlR* by exchanging the glutamic acid encoding-codon (GAA) to a nonsense mutation (TAA). The overlap fragments were subcloned between the EcoRI and BamHI sites of pMAD (1), yielding pIG1 and pIG6, respectively. pIT1, pIG1 and pIG6 were integrated in the *B. subtilis* chromosome by a single recombination event, forced by growth with the appropriate antibiotic at 42 ºC, for which the plasmid is non replicative. A second recombination event was promoted by growth at a permissive temperature without antibiotic, resulting in the restoration of the genotype or the clean introduction of the desired alteration. The detailed procedure for the generation of these clean alterations is described in detail by Arnaud *et al.*, 2004 (1). The in-frame Δ*frlO* construction resulted in the removal of a 525 bp region between amino acid-encoding positions 121 and 295, generating the *B. subtilis* strain ISN88. The 884 bp deletion of the S1254 region in the Δ*msmX*::*cat* background resulted in the construction of strain ISN51. The *frlRstop* mutation was introduced in a WT and in a Δ*msmX* background, yielding strains ISN71 and ISN72, respectively.

The construction of *B. subtilis* with 5’*frlP*-*lacZ* transcriptional and translational fusions in WT and *frlRstop* backgrounds was caried out by placing a *lacZ* gene copy of *E. coli* fused to the 5’-region of *frlP* using integrative plasmids. For these constructions pIG24, pIG25, pIG26, pIG27 and pIG28 were assembled. pIG24 is the backbone of the remaining vectors and was created by subcloning the *lacZ* region from pJM783 (2) into pJL3 (3) between KspAI and EcoRI sites. The *lacZ* gene was amplified by PCR with primers ARA826 and ARA1051 and treated with EcoRI and Bsp68I before cloning. For the construction of the remaining plasmids, the 5’ region of *frlP* was amplified by PCR from chromosomal DNA of *B. subtilis* 168T^+^ with primers ARA1052 and ARA1053, which contain additional EcoRI and SmaI/SalI restriction sites, respectively. The resulting fragments of approximately 378 bp were digested with EcoRI and SmaI for transcriptional fusions or with EcoRI and SalI for translational fusions. The products were cloned into pIG24 treated with the respective set of restriction enzymes, yielding plasmids pIG25 and pIG26 with transcriptional and translational fusions, respectively. These plasmids were used as templates for site-directed mutagenesis using mutagenic primers ARA929 and ARA930 to introduce the nonsense mutation *frlRstop* (described above), yielding pIG27 and pIG28, respectively. The fusions were introduced via transformation of *B. subtilis* 168T^+^ or *B. subtilis* ISN71 with the respective plasmids followed by a single crossover event, yielding strains ISN128 to ISN131 (Table S1).

All *B. subtilis* strains were transformed based on the protocol by Anagnostopoulos and Spizizen, 1960 (4). All modifications were confirmed by PCR and sequencing. Strains used and constructed for this work are summarized in Table 1.

**Bacterial Adenylate Cyclase Two-Hybrid (B2H) system: construction of expression vectors for protein-protein interaction studies**

To test protein-protein interactions between Nucleotide Binding Domains (NBDs) MsmX and FrlP and Transmembrane Domais (TMDs) FrlN and FrlM, the B2H System ((5); Euromedex) was used. Zip-Zip and MsmX-AraQ interactions were used as positive controls, and empty pKT25 and pUT28 were used as negative controls for interaction. The open reading frames (ORFs) of the ATPases were cloned in pUT18 vectors and the ORFs of permeases were cloned in pKT25/pKNT25 to generate in-frame fusions to the respective T18 or T25 fragments. We found that this combination of NBDs fused to T18 and TMDs fused to T25 was the one for which the interaction test was more efficient (unpublished work). The ORFs of *msmX* and *frlP* were PCR-amplified with primers ARA932 and ARA933 and ARA943 and ARA944, respectively, that introduced XbaI restriction sites on both 5’ and 3’ ends. These ORFs, treated with XbaI, were ligated to pUT18 digested with the same enzyme, yielding plasmids pLG55 and pJS1. Using primers containing additional restriction sites KpnI and BamHI, ARA945 and ARA951, *araQ* was amplified by PCR, treated with KpnI and BamHI and cloned into pKT25 similarly digested, yielding pLG64. Primers ARA1030 and ARA1031, containing XbaI restriction sites, were used for the PCR amplification of *frlN*, which was digested and cloned into pKNT25 similarly treated, leading to the construction of pIG20. The *frlM* ORF was amplified by PCR using primers ARA1033 and ARA1050, designed to introduce KpnI and XbaI restriction sites, digested with the same enzymes and cloned into pKT25, originating pIG23. All PCR amplifications described here were performed using chromosomal DNA of *B. subtilis* 168T^+^ as template. *E. coli* BTH101 was co-transformed with one pUT18 and one pKT25/pKNT25 derivatives, and the resulting co-transformants were selected in LA supplemented with ampicillin (100 μg mL^-1^), kanamycin (30 μg mL^-1^) and streptomycin (100 μg mL^-1^). These co-transformants were inoculated in LB with the respective antibiotics and IPTG (0.5 mM), grown aerobically at 30 ºC and protein-protein interaction was measured by β-galactosidase activity following Euromedex instructions, which was expressed in Units mg^-1^ dry weight bacteria.

### Construction of an expression vector for anti-FrlP antibody production

A His_6_-tag was added to the 3’-end of *frlP* via a single recombinational event into the chromosome to the *B. subtilis* strains for detection of FrlP (not shown). However, the doubling time of the strains in CSK supplemented with 0.1% (w/v) α- 1,5-arabinotriose as the sole carbon and energy source was severely affected. Therefore, it was imperative to construct an overexpression vector for heterologous FrlP production and posterior purification for synthesis of an anti-FrlP antibody. The FrlP coding sequence was amplified from *B. subtilis* WT with primers ARA837 and ARA960 containing unique restriction sites for NheI and HindIII, respectively, and cloned in pET28b(+) between the same restriction sites, creating the expression vector pIG9. The resulting His_6_-FrlP construction was heterologously expressed in *E. coli* BL21(DE3)pLysS. Cells were grown aerobically at 37 ºC in 125 mL LB medium with the appropriate antibiotic and the expression was induced at exponential growth phase by the addition of 0.1 mM IPTG and further grown by 16 h at 16 ºC and 150 rpm. Cells were collected by centrifugation at 9,000 x *g* for 10 min and the pellets were kept at -80 ºC until purification. All subsequent steps were performed on ice or at 4 ºC. The recovered pellet was resuspended in 10 mM PBS pH 7.4, 500 mM NaCl, 10 % (vol/vol) glycerol, 10 mM imidazole, 10 mM MgCl_2_, 5 mM 2-mercaptoethanol and 8 M Urea. Cell lysis was performed by sonication in the presence of 5 mU mL^-1^ Benzonase® Nuclease (Sigma), 1 mg mL^-1^ lysozyme and 10 mM phenylmethylsulfonyl fluoride (PMSF), and the lysis mixture was centrifuged at 13,000 x *g* for 1 h. The supernatant was passed through a 45 μm filter and loaded onto a 1-mL HisTrap column (GE Healthcare Life Sciences). Bounded proteins were eluted by discontinuous imidazole gradient; elution buffers have the same composition as lysis buffer, discriminated above, excluding imidazole concentration. The protein content was reinforced by repeating the purification procedure described above from an insoluble pellet fraction. Analysis of His_6_-FrlP production and purification was assessed by sodium dodecyl sulfate-polyacrylamide gel electrophoresis (SDS-PAGE) (12.5 %), using the Low Molecular Weight Marker from NZYTech as standard and Coomassie Blue reagent for protein staining. Fractions containing higher His_6_-FrlP content were concentrated using Amicon® Ultra Centrifugal Filter, 30 kDa MWCO (Milipore) and the obtained concentrated mixture was quantified using Bradford reagent (Bio-Rad Laboratories), using bovine serum albumin as standard. A 10 % SDS-PAGE hand cast gel loaded with the concentrated sample was run at constant electrical current (80 mA) for 50 min and the protein sample was visualized through the staining of a pilot lane. His_6_-FrlP was sent to Davids Biotechnologie GmbH (Germany) for polyclonal antibody synthesis.
